# Optical measurement of voltage sensing by endogenous ion channels

**DOI:** 10.1101/541805

**Authors:** Parashar Thapa, Robert Stewart, Rebecka J. Sepela, Oscar Vivas, Laxmi K. Parajuli, Mark Lillya, Sebastian Fletcher-Taylor, Bruce E. Cohen, Karen Zito, Jon T. Sack

## Abstract

A primary goal of molecular physiology is to understand how conformational changes of proteins affect the function of cells, tissues, and organisms. Here, we describe an imaging method for measuring the conformational changes of the voltage sensors of endogenous ion channel proteins within live tissue, without genetic modification. We synthesized GxTX-594, a variant of the peptidyl tarantula toxin guangxitoxin-1E, conjugated to a fluorophore optimal for two-photon excitation imaging through light-scattering tissue. GxTX-594 targets the voltage sensors of Kv2 proteins, which form potassium channels and plasma membrane–endoplasmic reticulum junctions. GxTX-594 dynamically labels Kv2 proteins on cell surfaces in response to voltage stimulation. To interpret dynamic changes in labeling intensity, we developed a statistical thermodynamic model that relates the conformational changes of Kv2 voltage sensors to labeling intensity. We used two-photon excitation imaging of rat brain slices to image Kv2 proteins in neurons. This imaging method enabled identification of conformational changes of endogenous Kv2 voltage sensors in tissue.

## INTRODUCTION

To move the field of voltage-sensitive physiology forward, we need new tools that indicate when and where voltage-sensitive conformational changes in endogenous proteins occur. Many classes of transmembrane proteins have been found to be voltage sensitive (Bezanilla 2008). One important class of voltage-sensitive proteins is the voltage gated ion channels. Electrophysiological techniques have enabled remarkably precise studies of the voltage-sensitivity of ionic conductances, primarily under reductionist experimental conditions where the channels have been removed from their native tissue. In addition to their canonical function as ion-conducting channels, voltage gated ion channel proteins have nonconducting functions which are independent of their ion conducting functions (Tanabe, Beam et al. 1988, Kaczmarek 2006). These nonconducting protein functions are largely inaccessible to study by electrophysiology and are more poorly understood. Novel approaches are needed to learn more about voltage-sensing in intact tissues and to unlock the mysterious realm of nonconducting voltage-sensitive physiology. Here, we present a new approach to image the conformational changes of endogenous voltage-sensitive proteins.

We have developed a protein imaging tool, an <u>E</u>ndogenous <u>V</u>oltage-sensor <u>A</u>ctivity <u>P</u>robe (EVAP), which dynamically labels living tissue to reveal the conformational status of endogenous Kv2 proteins with subcellular resolution. Kv2 proteins form voltage-gated K^+^ channels (Frech, VanDongen et al. 1989), bind endoplasmic reticulum proteins to form plasma membrane–endoplasmic reticulum junctions (Johnson, Leek et al. 2018, Kirmiz, Palacio et al. 2018), regulate a wide variety of physiological responses in tissues throughout the body (Bocksteins 2016), and integrate their response to voltage with many other cellular processes including phosphorylation (Murakoshi, Shi et al. 1997), sumoylation (Plant, Dowdell et al. 2011), oxidation (MacDonald, Salapatek et al. 2003), membrane lipid composition (Ramu, Xu et al. 2006), and auxiliary subunits (Gordon, Roepke et al. 2006, Peltola, Kuja-Panula et al. 2011).

Kv2 proteins are members of the voltage-gated cation channel superfamily. The voltage sensors of proteins in this superfamily are comprised of a bundle of four transmembrane helices termed S1-S4 (Long, Campbell et al. 2005, Long, Tao et al. 2007). The S4 helix contains positively charged arginine and lysine residues, gating charges, which respond to voltage changes by moving through the transmembrane electric field (Aggarwal and MacKinnon 1996, Seoh, Sigg et al. 1996, Islas and Sigworth 1999). When voltage sensor domains encounter a transmembrane voltage that is more negative on the inside of the cell membrane, voltage sensors are biased towards resting conformations, or down states, in which gating charges are localized intracellularly. When voltage becomes more positive, gating charges translate towards the extracellular side of the membrane, and voltage sensors are progressively biased towards up states, in a process of voltage activation (Armstrong and Bezanilla 1973, Zagotta, Hoshi et al. 1994, Tao, Lee et al. 2010, Xu, Li et al. 2019). Channel pore opening is distinct from, but coupled to, voltage sensor movement. In some voltage-gated ion channel proteins, voltage sensor movement is coupled to nonconducting protein functions (Tanabe, Beam et al. 1988, Kaczmarek 2006). To study the functional outputs of voltage sensors it is essential to measure voltage sensor activation itself. Conformational changes of voltage sensors have been detected with electrophysiological measurements of gating currents (Armstrong and Bezanilla 1973, Schneider and Chandler 1973, Bezanilla 2018) or by optical measurements from fluorophores covalently attached to voltage sensors by genetic encoding (Lin and Schnitzer 2016) or by chemical engineering (Zhang, Zheng et al. 2015). However, experimental limitations prevent these existing techniques from measuring conformational changes of voltage sensors of most endogenous proteins: gating currents can only be measured when the proteins are expressed at high density in a voltage-clamped membrane; engineered proteins differ from endogenous channels; and introducing point mutations and conjugated fluorophores into voltage sensors alters structure and function. Here, we develop a different strategy, EVAPs that dynamically label endogenous voltage sensors of Kv2 proteins to reveal conformational states.

To image where the voltage sensors of Kv2 proteins adopt a specific resting conformation in tissue, we exploit the conformation-selective binding of the tarantula peptide guangxitoxin-1E (GxTX), which can be conjugated to fluorophores to report Kv2 conformational changes (Tilley, Eum et al. 2014). Here, we synthesize GxTX-594, a Ser13Cys GxTX variant conjugated to Alexa Fluor 594, a fluorophore compatible with two-photon excitation imaging through light-scattering tissue. This EVAP equilibrates with Kv2 proteins on the time scale of seconds, revealing the probability (averaged over time) that unbound voltage sensors are resting or active. We develop a method to calculate the average conformational status of unlabeled Kv2 proteins from images of GxTX-594 fluorescence. We deploy the GxTX-594 probe in brain slices and image voltage-sensitive fluorescence changes that reveal conformational changes of endogenous neuronal Kv2 proteins. This EVAP approach provides an imaging technique to study conformational changes of voltage-sensitive proteins in samples that have not (or cannot) be genetically modified.

## MATERIALS AND METHODS

### GxTX-594 synthesis

The Ser13Cys GxTX peptide was synthesized as described (Tilley, Eum et al. 2014). Methionine 35 of GxTX was replaced by the oxidation-resistant noncanonical amino acid norleucine and serine 13 was replaced with cysteine to create a spinster thiol. Ser13Cys GxTX was labeled with a Texas red derivative (Alexa Fluor 594 C_5_ maleimide, Thermo Fisher Scientific, Catalog # 10256) to form GxTX-594. Ser13Cys GxTX lyophilisate was brought to 560 μM in 50% ACN + 1 mM Na_2_EDTA. 2.4 μL of 1M Tris (pH 6.8 with HCl), 4 μL of 10 mM of Alexa Fluor 594 C_5_ maleimide in DMSO and 17.9 μL of 560 μM Ser13Cys GxTX were added for a final solution of 100 mM Tris, 1.6 mM Alexa Fluor 594 C_5_ maleimide and 0.4 mM GxTX in 24 μL of reaction solution. Reactants were combined in a 1.5 mL low protein binding polypropylene tube (LoBind, Eppendorf, Catalog # 022431081) and mixed at 1000 RPM, 20°C for 4 hours (Thermomixer 5355 R, Eppendorf). After incubation, the tube was centrifuged at 845 RCF for 10 minutes at room temperature. A purple pellet was observed post-centrifugation. The supernatant was transferred to a fresh tube and centrifuged at 845 RCF for 10 minutes. After this 2^nd^ centrifugation, no visible pellet was seen. The supernatant was injected onto a reverse-phase HPLC C_18_ column (Biobasic-4.6mm RP-C_18_ 5 µm, Thermo Fisher Scientific, Catalog # 2105-154630) equilibrated in 20% ACN, 0.1% TFA at 1 mL/min and eluted with a protocol holding in 20% ACN for 2 minutes, increasing to 30% ACN over 1 minute, then increasing ACN at 0.31% per minute. HPLC effluent was monitored by fluorescence and an absorbance array detector. 1 mL fractions were pooled based on fluorescence (280 nm excitation, 350 nm emission) and absorbance (214 nm, 280 nm, and 594 nm). GxTX-594 peptide-fluorophore conjugate eluted at approximately 35% ACN, and mass was confirmed by mass spectrometry using a Bruker Ultra Flextreme MALDI-TOF/TOF (Fig 1). Samples for identification from HPLC eluant were mixed 1:1 in an aqueous solution of 25% MeOH and 0.05% TFA saturated with alpha-cyano-4-hydrocinnamic acid, pipetted onto a ground-steel plate, dried under vacuum, and ionized with 60-80% laser power. Molecular species were detected using a reflector mode protocol and quantitated using Bruker Daltonics flexAnalysis 3.4. Lyophilizate containing GxTX-594 conjugation product was dissolved in CE buffer (defined below) and stored at −80 °C. GxTX-594 concentration was determined by 280 nm absorbance using a calculated extinction coefficient of 18,900 A.U. M^-1^ cm^-1^.

**Figure 1:**
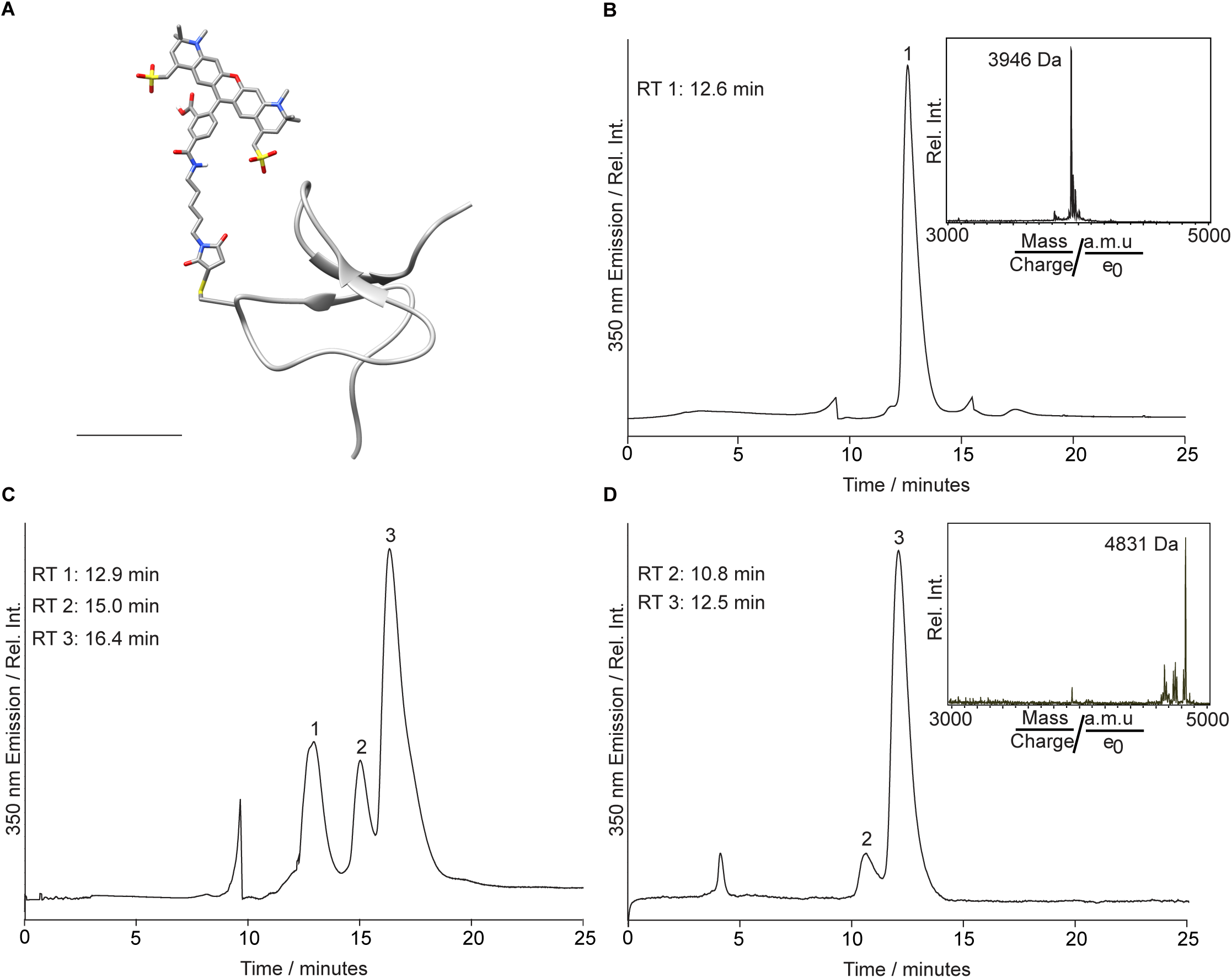
Synthesis of GxTX-594. A. Molecular model of GxTX-594. Scale bar is 10 angstroms. Backbone of GxTX peptide depicted with ribbon. Cys13(maleimide-Alexa 594) depicted with CPK atoms. B. HPLC chromatogram of Ser13Cys GxTX. Gradient described in *Materials and Methods - GxTX Synthesis*. Ser13Cys GxTX eluted at 12.6 minutes, peak 1, which corresponds to 33% acetonitrile. MALDI-TOF MS profile of peak 1 (Inset). Rel. Int. is Relative Intensity. C. HPLC chromatogram of GxTX-594 conjugation reaction between Alexa Fluor 594-maleimide and Ser13Cys GxTX. Peak 1 is Ser13Cys GxTX (Retention time: 12.8 minutes, 33% acetonitrile), peak 2 is a minor product from conjugation, and peak 3 is GxTX-594, the major product from conjugation (Retention time: 16.4 minutes, 35% acetonitrile). The fractions corresponding to peak 3 were combined. D. HPLC chromatogram of the combined peak 3 fractions from panel C, a GxTX-594 preparation used in this study. 2 μL of 13.1 µM GxTX-594 diluted in 200 μL of 0.1 % TFA was injected. MALDI-TOF MS profile of the combined peak 3 fractions (Inset).

### CHO cell methods

#### CHO cell culture and transfection

The CHO-K1 cell line (ATCC) and a subclone transfected with a tetracycline-inducible rat Kv2.1 construct (Kv2.1-CHO) (Trapani and Korn 2003), were cultured as described previously (Tilley, Eum et al. 2014). To induce Kv2.1 expression in Kv2.1-TREx-CHO cells, 1 µg/mL minocycline (Enzo Life Sciences, Catalog # ALX-380-109-M050), prepared in 70% ethanol at 2 mg/mL, was added to the maintenance media to induce K^+^ expression. Minocycline was added 40-48 hours before imaging and voltage clamp fluorometry experiments. Minocycline was added 1-2 hours before whole cell ionic current recordings, to limit K^+^ conductance such that voltage clamp could be maintained. Transfections were achieved with Lipofectamine 2000 (Life Technologies, Catalog # 1668027). 1.1 μL of Lipofectamine was diluted, mixed, and incubated in 110 μL of Opti-MEM (Gibco, Product Number 31985062, Lot Number 1917064) in a 1:100 ratio for 5 minutes at room temperature. Concurrently, 1 μg of plasmid DNA and 110 μL of Opti-MEM were mixed in the same fashion. DNA and Lipofectamine 2000 mixtures were combined, titurated, and let sit for 20 minutes. Then, the transfection cocktail mixture was added to a 35 mm cell culture dish of CHO cells at approximately 40% confluency and allowed to settle at 37°C 5% CO_2_ for 4-6 hours before the media was replaced. Rat Kv2.1-GFP (Antonucci, Lim et al. 2001), rat Kv2.2-GFP (Kirmiz, Vierra *et al*. 2018), rat Kv1.5-GFP (Li, Guo et al. 2001), rat Kv4.2-GFP (Shibata, Misonou et al. 2003), mouse BK-GFP (Trimmer Lab, UC Davis), rat Kv2.1 S586A-GFP (Kirmiz, Palacio et al. 2018), rat AMIGO-YFP (Bishop, Guan et al. 2015), rat KvBeta2 (Shibata, Misonou et al. 2003) and rat KCHiP2 (An, Bowlby et al. 2000) plasmids were all kind gifts from James Trimmer, University of California, Davis. Identities of constructs were confirmed by sequencing from their CMV promoter.

#### Cell transfection for electrophysiology

Prior to transfection, cells were plated at 40% confluency in culture media free of selection agents. Transfections were achieved with Lipofectamine 2000 (Life Technologies, Catalog # 1668027) following the manufacturer’s protocol. Transfection mixes were prepared in Opti-MEM (Life Technologies, Product Number 31985062) and included 1 μg of peGFP. Cells were incubated in the transfection cocktail for 6-8 hours before being returned to regular growth media. Cells were given 40-48 hours recovery following transfection before being used for experiments.

#### Confocal and Airy disk imaging

Confocal images were obtained with an inverted confocal system (Zeiss LSM 880 410900-247-075) run by ZEN black v2.1. A 63x/1.40 NA Oil DIC objective (Zeiss 420782-9900-799) was used for most imaging experiments; a 63x/1.2 NA Water DIC objective (Zeiss 441777-9970-000) was used for voltage clamp fluorometry experiments. GFP and YFP were excited with the 488 nm line from an argon laser (3.2 mW at installation) powered at 0.5% unless otherwise noted. GxTX-594 was excited with 594 nm helium-neon laser (0.6 mW at installation) powered at 10% unless otherwise noted. WGA-405 was excited with a 405 nm diode laser, (3.5 mW at installation) powered at 1% unless otherwise noted. Temperature inside the microscope housing was 27-30 °C.

In confocal imaging mode, fluorescence was collected with the microscope’s 32-detector gallium arsenide phosphide detector array arranged with a diffraction grating to measure 400-700 nm emissions in 9.6 nm bins. Emission bands were 495-550 nm for GFP and YFP, 605-700 nm for GxTX-594, and 420-480 nm for WGA-405. Point spread functions were calculated using ZEN black software using emissions from 0.1 μm fluorescent microspheres, prepared on a slide according to manufacturer’s instructions (Thermo Fisher Scientific, Catalog # T7279). The point spread functions for confocal images with the 63x/1.40 Oil DIC objective in the X-Y direction were 228 nm (488 nm excitation) and 316 nm (594 nm excitation).

In Airy disk imaging mode, fluorescence was collected with the microscope’s 32-detector gallium arsenide phosphide detector array arranged in a concentric hexagonal pattern (Zeiss Airyscan 410900-2058-580). After deconvolution, the point spread functions for the 63x/1.40 NA objective with 488 nm excitation was 124 nm in X-Y and 216 nm in Z; with 594 nm excitation 168 nm in X-Y and 212 nm in Z. For the 63x/1.2 NA objective, the point spread function with 488 nm excitation was 187 nm in X-Y and 214nm in Z; the point spread function with 594 nm excitation was 210 nm in X-Y and 213 nm in Z.

Unless stated otherwise, cells were plated in uncoated 35 mm dishes with a 7 mm inset No. 1.5 coverslip (MatTek, Catalog # P35G-1.5-20-C). The CHO cell external (CE) solution used for imaging and electrophysiology contained (in mM): 3.5 KCl, 155 NaCl, 10 HEPES, 1.5 CaCl_2_, 1 MgCl_2_, adjusted to pH 7.4 with NaOH. Measured osmolarity was 315 mOsm. When noted, solution was supplemented with either 10 mM glucose (CEG) or 10 mM glucose and 1% BSA (CEGB).

For time-lapse, GxTX-594 dose-response experiments, Kv2.1-CHO cells were plated onto 22 x 22 mm No. 1.5H coverglass (Deckglaser). Prior to imaging, cell maintenance media was removed and replaced with CEGB, then coverslip was mounted on an imaging chamber (Warner Instrument, Catalog # RC-24E) with vacuum grease. During time-lapse imaging, indicated concentrations of GxTX-594 were applied for 10 minutes followed by manual wash-out with CEGB solution. 15 minutes after wash-out, the next GxTX-594 concentration was added. Solutions were added to the imaging chamber perfusion via a syringe at a flow rate of ∼1 mL / 10 sec. Images were taken every 5 sec. Laser power was set to 0.5% for the 488 nm laser and 1.5% for the 594 nm laser. For colocalization experiments with GFP-tagged proteins, cells were incubated in 100 nM GxTX-594 for 5 minutes and then washed with 1 mL of CEGB three times before imaging.

#### Whole cell voltage clamp for CHO cell imaging

Kv2.1-TREx-CHO cells plated in glass-bottom 35 mm dishes were washed with CEGB, placed in the microscope, and then incubated in 100 µL of 100 nM GxTX594 for 5 minutes to label cells. Before patch clamp, the solution was diluted with 1 mL of CEG for a working concentration of 9 nM GxTX594 during experiments. Cells with obvious GxTX-594 surface staining were voltage-clamped in whole-cell mode with an EPC-10 patch clamp amplifier (HEKA) run by Patchmaster software, v2×90.2 (HEKA). The patch pipette contained a potassium-deficient Cs^+^ internal pipette solution to limit outward current and reduce voltage error: 70 mM CsCl, 50 mM CsF, 35mM NaCl, 1 mM EGTA, 10 mM HEPES, brought to pH 7.4 with CsOH. Osmolarity was 310 mOsm. The liquid junction potential was calculated to be 3.5 mV and was not corrected. Borosilicate glass pipettes (Sutter Instruments, Catalog # BF 150-110-10HP) were pulled with blunt tips to resistances less than 3.0 MΩ in these solutions. Cells were held at −80 mV (unless noted otherwise) and stepped to indicated voltages. The voltage step stimulus was maintained until any observed change in fluorescence was complete. For stimulus-frequency dependence experiments, cells were given 2 ms steps to 40 mV at stated frequencies (0.02, 5, 10, 20, 50, 100, 150, or 200 Hz). Images for voltage clamp fluorometry were taken in Airy disk imaging mode with the settings described above.

#### K^+^ channel-GFP ionic current recordings

Whole-cell voltage clamp was used to measure currents from CHO cells expressing Kv2.1-GFP, Kv2.2-GFP, Kv1.5-GFP, Kv4.2-GFP, BK-GFP or GFP that was transfected as described above. Cells were plated on glass bottom dishes. Cells were held at −80 mV, then 100 ms voltage steps were delivered ranging from −80 mV to +80 mV in +5 mV increments. Pulses were repeated every 2 seconds. The external (bath) solution contained CE solution. The internal (pipette) solution contained (in mM): 35 KOH, 70 KCl, 50 KF, 50 HEPES, 5 EGTA adjusted to pH 7.2 with KOH. Liquid junction potential was calculated to be 7.8 mV and was not corrected. Borosilicate glass pipettes (Sutter Instruments, Cat # BF150-110-10HP) were pulled into pipettes with resistance less than 3 MΩ for patch clamp recording. Recordings were at room temperature (22–24 °C). Voltage clamp was achieved with an Axon Axopatch 200B amplifier (MDS Analytical Technologies) run by Patchmaster software, v2×90.2 (HEKA). Holding potential was −80 mV. Capacitance and Ohmic leak were subtracted using a P/5 protocol. Recordings were low pass filtered at 10 kHz and digitized at 100 kHz. Voltage clamp data were analyzed and plotted with Igor Pro 7 or 8 (Wavemetrics). Current amplitudes at each voltage were the average from 0.19-0.20 s after voltage step. As the experiments plotted in Figure 6 Supplement 1 A were merely to confirm functional expression of ion channels at the cell surface, series resistance compensation was not used, and substantial cell voltage errors are predicted during these experiments.

#### Kv2.1 ionic current recordings

Prior to patching, Kv2.1-CHO cells were washed in divalent-free PBS, then harvested in Versene (Gibco, Cat # 15040066). Cells were scraped and transferred to a polypropylene tube, pelleted and washed three times at 1,000 g for 2 minutes and then resuspended in the same external solution as used in the recording chamber bath. Cells were rotated in a polypropylene tube at room temperature (22-24°C) until use. Cells were then pipetted into a 50 uL recording chamber (Warner Instruments, RC-24N) prefilled with external solution and allowed to settle for 5 or more minutes. After adhering to the bottom of the glass recording chamber, cells were thoroughly rinsed with external solution using a gravity-driven perfusion system. Cells showing uniform intracellular GFP expression of intermediate intensity were selected for patching.

Voltage clamp was achieved with a patch clamp amplifier (Axon Instruments Axopatch 200B) run by Patchmaster software (HEKA). Borosilicate glass pipettes (Sutter, BF150-110-7.5HP) were pulled with blunt tips, coated with silicone elastomer (Dow Corning Sylgard 184), heat cured, and tip fire-polished to resistances less than 4MΩ. Capacitance and Ohmic leak were subtracted using a P/5 protocol. Recordings were low-pass filtered at 10kHz using the amplifier’s built-in Bessel and digitized at 100kHz.

For whole cell ionic currents measurements in Kv2.1-CHO cells, the external patching solution contained (in mM): 3.5 KCl, 155 NaCl, 10 HEPES, 1.5 CaCl2, 1 MgCl2, adjusted to pH 7.4 with NaOH. The internal (pipette) solution contained (in mM): 70 KCl, 5 EGTA, 50 HEPES, 50 KF, and 35 KOH, adjusted to pH 7.4 with KOH. The osmolarity for the external solution was 315 mOsm, and 310 mOsm for the internal solution measured by a vapor presume osmometer. Following establishment of the whole-cell seal, ionic K+ current recordings were taken in the presence of a vehicle which consisted of 100 nM TTX, 10 mM glucose, and 0.1% BSA prepared in external solution. Cells were held at −100 mV with channel activation steps ranging from −80 mV to +120 mV in increments of +5 mV (100 ms) before being returned to 0 mV (100 ms) to record tail currents. The inter-sweep interval was 2 s. To determine the bioactivity of GxTX-594, Kv2.1 ionic currents were recorded once more, 5 minutes following the wash-in of bath solution also containing 100 nM GxTX-594. Wash-ins were carried out while holding at −100 mV; during some wash-ins, the membrane potential was pulsed to 0 mV to gauge the time course of binding and channel inhibition. To exchange solution, 100 uL was washed through the chamber and removed distally through vacuum tubing to maintain constant bath fluid level.

#### Ionic current analysis

The average current in the 100 ms prior to step was used to zero-subtract the recording. Outward current taken as the mean value between 90-100 ms of the channel activation step and used to calculate and correct for series resistance induced voltage error. Tail current values were derived from the mean value between 0.2-1.2 ms of the 0 mV tail current step. Tail current was normalized by the mean activation step current from 50-80 mV and plotted against the estimated membrane potential, which had been corrected for voltage-error and the calculated liquid junction potential of 8.5 mV. These tail GV plots were fit with a 4^th^ power Boltzmann (Sack, Aldrich et al. 2004) and the fit parameters were used for statistical analysis.

### Brain slice methods

#### Hippocampal slice culture preparation and transfection

All experimental procedures were approved by the University of California Davis Institutional Animal Care and Use Committee and were performed in strict accordance with the Guide for the Care and Use of Laboratory Animals of the NIH. Animals were maintained under standard light– dark cycles and had ad libitum access to food and water. Organotypic hippocampal slice cultures were prepared from postnatal day 5-7 rats, as described (Stoppini, Buchs et al. 1991) and detailed in a video protocol (Opitz-Araya and Barria 2011). DIV15-30 neurons were transfected 2-6 days before imaging via biolistic gene transfer (160 psi, Helios gene gun, Bio-Rad), as described in a detailed video protocol (Woods and Zito 2008). 10 µg of plasmid was coated to 6-8 mg of 1.6 µm gold beads.

#### Two-photon excitation slice imaging

Image stacks (512 x 512 pixels, 1 mm Z-steps; 0.035 μm/pixel) were acquired using a custom two-photon excitation microscope (Olympus LUMPLFLN 60XW/IR2 objective, 60x, 1.0 NA) with two pulsed Ti:sapphire lasers (Mai Tai: Spectra Physics) tuned to 810 nm (for GxTX-594 imaging) and 930 nm (for GFP imaging) and controlled with ScanImage (Pologruto, Sabatini et al. 2003). After identifying a neuron expressing Kv2.1-GFP, perfusion was stopped and GxTX-594 was added to the static bath solution to a final concentration of 100 nM. After five minutes incubation, perfusion was restarted, leading to wash-out of GxTX-594 from the slice bath. Red and green photons (Chroma: 565dcxr, BG-22 glass, HQ607/45) emitted from the sample were collected with two sets of photomultiplier tubes (Hamamatsu R3896).

#### Whole cell voltage clamp for brain slice imaging

Organotypic hippocampal slice cultures (DIV 6-7, not transfected) were transferred to an imaging chamber with recirculating ACSF maintained at 30°C. To hold the slice to the bottom of the chamber, a horseshoe-shaped piece of gold wire was used to weight the membrane holding the slice. ACSF contained (in mM) 127 NaCl, 25 NaHCO_3_, 25 D-glucose, 2.5 KCl, 1.25 NaH_2_PO_4_, 1 MgCl2, 2 CaCl_2_, 200 nM tetrodotoxin, pH 7.3 and aerated with 95% O_2_/ 5% CO_2_ (∼310mOsm). 4 mL of 100 nM GxTX-594 in ACSF were used in the circulating bath to allow the toxin to reach the slice and reach the desired concentration of 100 nM throughout the circulating bath. Images were acquired beginning 3 minutes after GxTX-594 was added.

Apparent CA1 neurons with GxTX-594 labeling in a Kv2-like pattern were selected for whole-cell patch clamp. Voltage clamp was achieved using an Axopatch 200B amplifier (Axon Instruments) controlled with custom software written in Matlab (MathWorks, Inc.). Patch pipettes (5-7 MΩ) were filled with intracellular solution containing (in mM): 135 Cs-methanesulfonate, 10 Na_2_-phosphocreatine, 3 Na-L-ascorbate, 4 NaCl, 10 HEPES, 4 MgCl_2_, 4 Na_2_ATP, 0.4 NaGTP, pH 7.2. Neurons were clamped at −70 mV. Input resistance and holding current were monitored throughout the experiment. Cells were excluded if the pipette series resistance was higher than 25 MΩ or if the holding current exceeded −100 pA. To activate Kv2 channels, a 50 s depolarizing step from −70 mV to 0 mV was given.

### Image analysis

Fluorescence images were analyzed using ImageJ 1.52n software (Schneider, Rasband et al. 2012). Regions of interest (ROIs) encompassed the entire fluorescent region of an individual cell or neuron, unless mentioned otherwise. ROIs were drawn manually. Analysis of images was conducted independently by multiple researchers who produced similar results, but analysis was not conducted in a blinded nor randomized fashion. Fluorescence intensity (*F)* was background subtracted using the mean *F* of a region that did not contain cells. In experiments with CHO cells where the bath solution contained 9 nM GxTX-594, the apparent surface membrane of most cells (40/47) which lack Kv2.1 protein had lower *F* than the background region that did not contain cells (Fig 4 A, horizontal black dotted line) indicating that the background *F* was overestimated. Based on the mean *F* from the apparent surface membrane of cells lacking Kv2.1 protein, the background was overestimated by 24%. The signal from the surface membrane of cells stably transfected with rat Kv2.1 (Kv2.1-CHO) was, on average, 18x higher than regions that did not contain cells, making the error generated by our overestimation of the background about 1%. As the error produced by this background subtraction method was relatively small, it was not corrected. For *F/F_init_* normalization, *F_init_* was the mean fluorescence preceding the indicated voltage stimuli, or the max observed intensity in dose response experiments. Further details of specific normalization and background subtraction procedures are provided in figure legends. Time dependence of fluorescence intensity was fit with a monoexponential decay (Eq. 1). For colocalization analyses the Pearson coefficient was calculated using the JACoP plugin (Bolte and Cordelieres 2006). Colocalization analyses were conducted within ROIs defining individual cells. Plotting and curve fitting was performed with Igor Pro 7 or 8 (Wavemetrics), which performs nonlinear least-squares fits using a Levenberg-Marquardt algorithm. Sample sizes of n ≥ 3 were selected to confirm reproducibility. Sample sizes of n ≥ 6 were selected to power nonparametric statistical comparisons to discern p < 0.01. Error values from individual curve fittings are standard deviations. All other errors, including error bars, indicate standard errors. Arithmetic means are reported for intensity measurements and correlation coefficients. Geometric means are reported for time constants and rates. As the distributions underlying variability in results are unknown, nonparametric statistical comparisons were conducted with Mann-Whitney U tests and two-tailed p values reported individually if p > 0.0001.

### Model calculations

Predictions of the Scheme I and II models were calculated using Microsoft Excel. A spreadsheet containing model calculations that can be used to generate model prediction is included as a supplement (Model Calculation Supplement).

## RESULTS

### GxTX-594 labels Kv2 proteins

To monitor activation of Kv2 proteins in tissue slices, we synthesized an EVAP compatible with two-photon imaging. We previously presented an EVAP that was a synthetic derivative of GxTX conjugated to a DyLight 550 fluorophore (GxTX-550) (Tilley, Eum et al. 2014). DyLight 550 has poor two-photon excitation properties, and for this study it was replaced with Alexa Fluor 594, a persulfonated Texas Red analog with a large two-photon excitation cross-section and ample spectral separation from green fluorescent protein (GFP), making it well-suited for multiplexed, two-photon excitation imaging experiments (Zito, Knott et al. 2004).

We used solid-phase peptide synthesis to generate a variant of GxTX, an amphiphilic 36-amino acid cystine knot peptide. We synthesized the same peptide used for GxTX-550, Ser13Cys GxTX, where a free thiolate side chain of cysteine 13 is predicted to extend into extracellular solution when the peptide is bound to a voltage sensor (Tilley, Eum et al. 2014). GxTX-1E folds by formation of 3 internal disulfides, and cysteine 13 was differentially protected during oxidative refolding to direct chemoselective conjugation. Following refolding and thiol deprotection, Alexa Fluor 594 C_5_ maleimide was condensed with the free thiol, and Ser13Cys (Alexa Fluor 594) GxTX-1E (called “GxTX-594” throughout) was purified (Fig 1).

We performed a dose–response experiment to determine the concentration range where GxTX-594 effectively labels Kv2.1 protein. For these experiments, Kv2.1 was expressed in a CHO-K1 cell line stably transfected with rat Kv2.1 under control of a tetracycline-inducible promoter (Kv2.1-CHO) (Trapani and Korn 2003). Kv2.1 expression was induced with minocycline 2 days prior to experiments, such that all Kv2.1-CHO cells expressed Kv2.1. The K_d_ of our previously presented EVAP, GxTX-550, was 30.0 ± 3.9 nM which was calculated from electrophysiology at a holding potential of −100 mV (Tilley, Eum et al. 2014) and we guessed that 33 times this K_d_, 1000 nM, would be a sufficient upper bound for dose-response experiments with the related GxTX-594 peptide. We performed three tenfold serial dilutions of 1000 nM GxTX-594 to generate the range of concentrations used for this dose–response and applied each concentration of GxTX-594 to Kv2.1-CHO cells for 15 minutes followed by 15 minutes of washout before the next concentration of GxTX-594 was applied (Fig 2). Due to the incomplete equilibration of labeling at 1 and 10 nM, the true K_d_ value for GxTX-594 is expected to be somewhat lower, and our measure is expected to be an overestimate of K_d_. A Langmuir binding isotherm was fit to fluorescence intensity immediately after washout of GxTX-594 from the extracellular solution and resulted in a K_d_ of 26.9 ± 8.3 nM (Fig 2 C). This suggests that GxTX-594 has a similar or higher affinity for resting voltage sensors than GxTX-550. The near saturation of the fluorescence dose-response with 100 nM GxTX-594 and rate of fluorescence equilibration (Fig. 2 D) suggested that incubation with 100 nM GxTX-594 for 5 minutes should be sufficient for near maximal labeling of Kv2.1-CHO cells.

**Figure 2:**
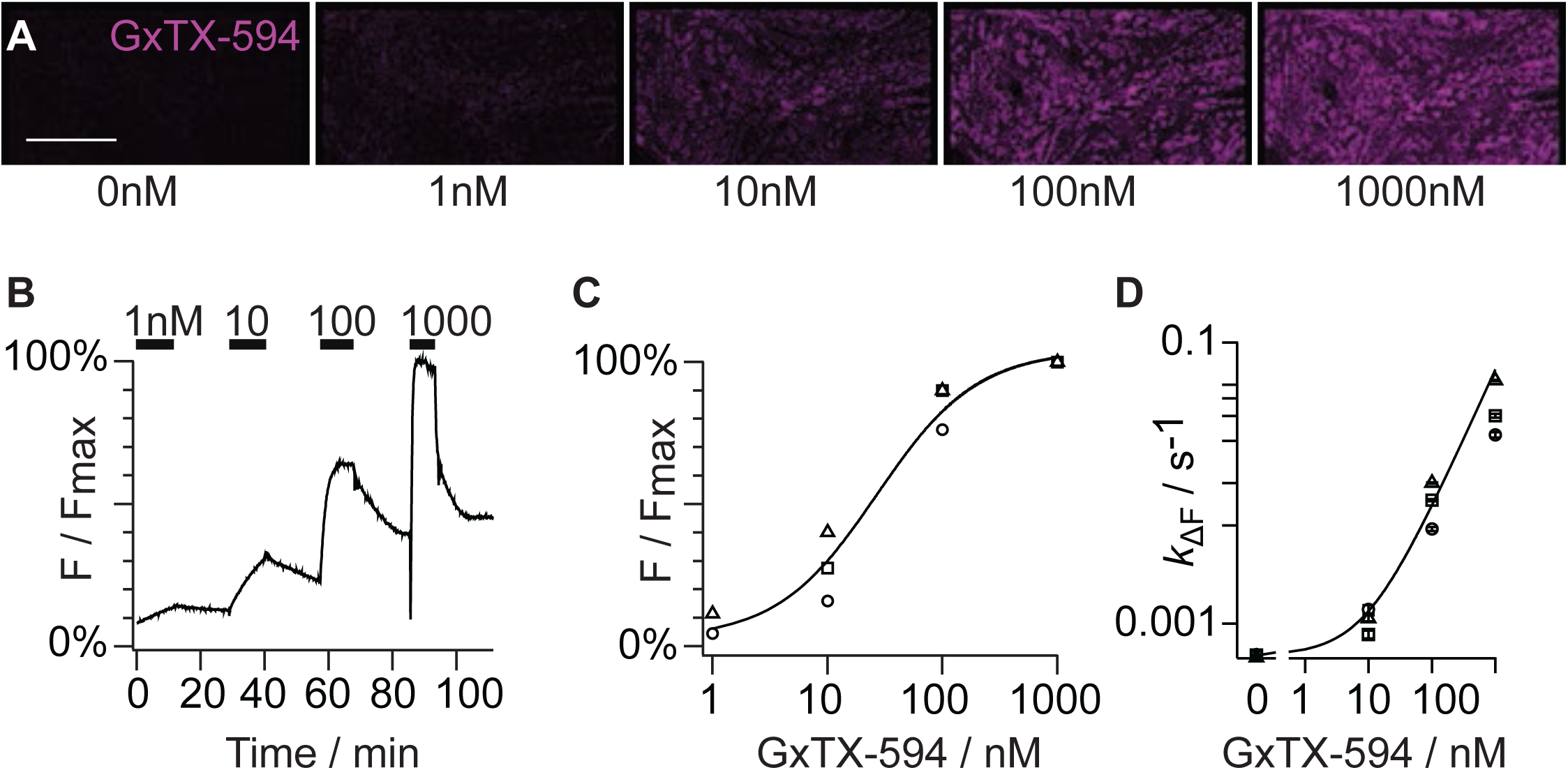
Dose-response characteristics of GxTX-594 labeling. A. Fluorescence from Kv2.1-CHO cells incubated in indicated concentrations of GxTX-594 for 15 minutes then washed out before imaging. Imaging plane was near the glass-adhered cell surface. Fluorescence shown corresponds GxTX-594 (Magenta). Scale bar is 20 µm. B. Time-lapse fluorescence intensity from the dose-response experiment shown in A. Fluorescence was not background-subtracted. F_max_ is intensity while cells were incubated in 1000 nM GxTX-594. C. Relative fluorescence intensity of GxTX-594 that remains on cells immediately after wash-out of indicated concentrations of GxTX-594 from the extracellular solution. Symbols correspond to each of 3 experiments. Black line is the fit of a Langmuir isotherm to obtain a dissociation constant (*K_d_*) for GxTX-594; *K_d_* = 26.9 nM ± 8.3. D. Labeling or unlabeling kinetics for GxTX-594 at indicated concentrations. Symbols correspond to each of 3 experiments. The *k*_ΔF_ from each point was obtained by a monoexponential fit (Eq. 1). Error bars represent the standard deviation of. Data were fit with Equation 4 (k_on_ = 6.372 x 10^-5^ ± 0.049 x 10^-5^ nM^-1^ s^-1^; k_off_ = 5.929 x 10^-4^ ± 0.043 x 10^-4^ s^-1^).

We assessed whether GxTX-594 fluorescence on Kv2.1-CHO cells is due to binding Kv2 proteins. Mammals have two pore-forming Kv2 proteins, Kv2.1 and Kv2.2, both of which are expressed in neurons throughout the brain (Trimmer 1991, Kihira, Hermanstyne et al. 2010, Bishop, Guan et al. 2015). CHO-K1 cells were transfected with either a rat Kv2.1-GFP or Kv2.2-GFP construct. We confirmed surface expression of these constructs by whole cell patch clamp electrophysiology. Transfection with either Kv2.1-GFP or Kv2.2-GFP yielded delayed rectifier K^+^ currents, consistent with Kv2 channels (Fig Supplement 1 A). The CHO-K1 cell line does not have endogenous K^+^ currents (Gamper, Stockand et al. 2005). Two days post transfection, Kv2.1-GFP or Kv2.2-GFP fluorescence intensity was non-uniform as seen previously for Kv2.1-GFP (Tilley, Eum et al. 2014), with distinct subcellular regions of high density separated by regions of much lower density (Fig 3 A). In neurons and other mammalian cell lines similar non-uniformities have been referred to as “clusters,” a term we adopt here to refer to high density regions of Kv2 channels. In HEK293 cells and neurons, Kv2 clusters are sites where VAP proteins in the endoplasmic reticulum bind Kv2 proteins in the plasma membrane to form endoplasmic reticulum–plasma membrane junctions (Johnson, Leek et al. 2018, Kirmiz, Palacio et al. 2018). A similar mechanism may underly the clustering of Kv2 proteins in CHO cells.

**Figure 3:**
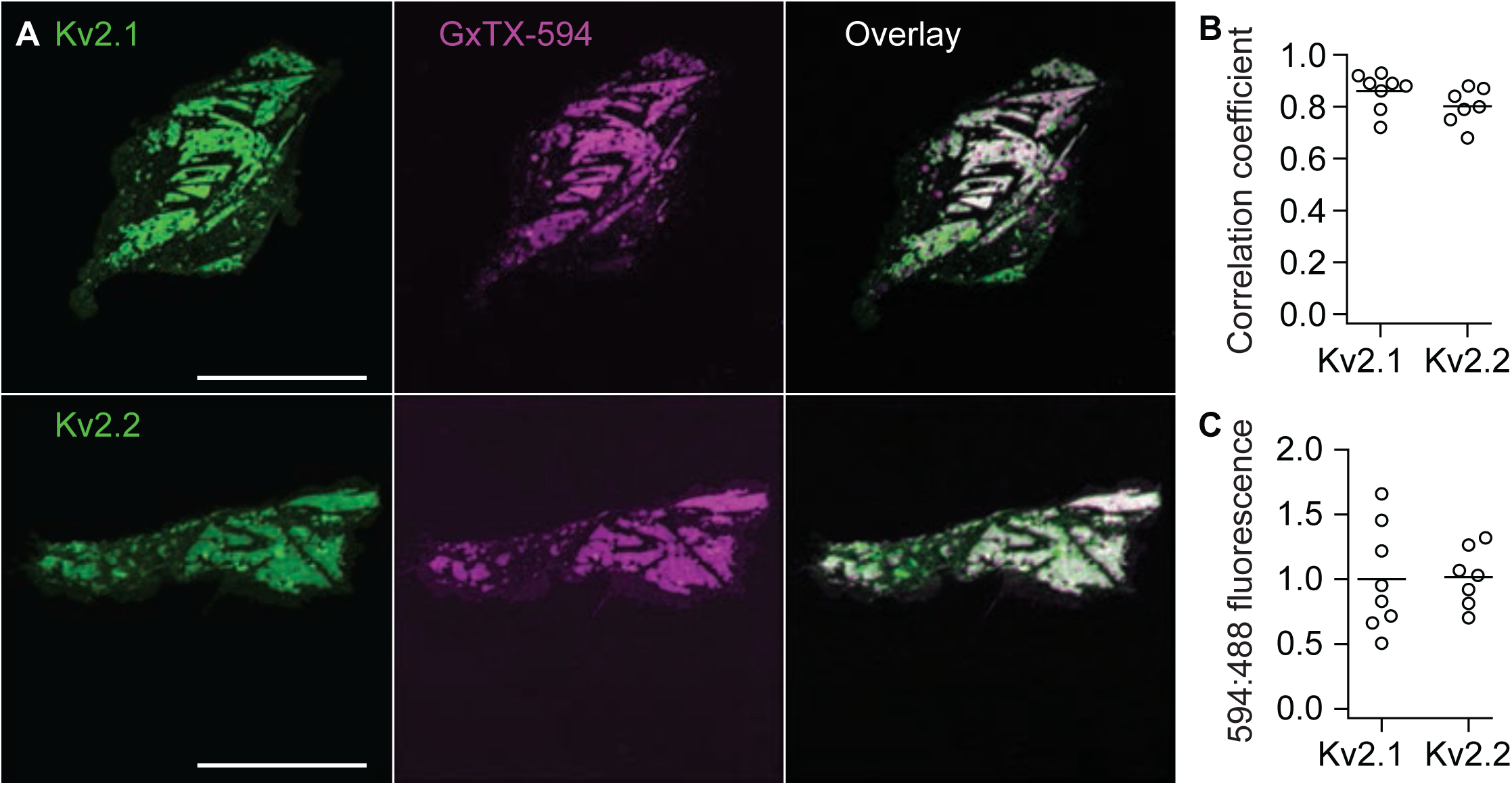
GxTX-594 colocalizes with Kv2-GFP. A. Fluorescence from CHO cells transfected with Kv2.1-GFP (Top) or Kv2.2-GFP (Bottom) and labeled with GxTX-594. Optical sections were near the glass-adhered cell surface. Cells were incubated with 100 nM GxTX-594 for 5 minutes and rinsed before imaging. Fluorescence shown corresponds to emission of GFP (Left), GxTX-594 (Middle), or an overlay of GFP and GxTX-594 (Right). Scale bars are 20 µm. B. Pearson correlation coefficient for GxTX-594 colocalization with Kv2.1-GFP or Kv2.2-GFP. In panels B and C pixel intensities were background subtracted before analyses by subtracting the average fluorescence of a background ROI that did not contain cells, from the ROI containing the cell. All data from the same application of GxTX-594. Circles represent measurements from individual cells. C. Ratio of fluorescence intensity when excited at 594 nm versus 488 nm from cells expressing Kv2.1-GFP or Kv2.2-GFP. The consistent ratio between Kv2.1 and Kv2.2 indicates that GxTX-594 labels the two proteins equivalently.

To determine the degree of association between GxTX-594 and Kv2 proteins in CHO cells we calculated the Pearson Correlation Coefficient (PCC) between the fluorescence intensity from GxTX-594 and Kv2-GFP proteins (Dunn, Kamocka et al. 2011). A PCC value of 0 signifies uncorrelated pixels between two images and a PCC value of 1 signifies complete correlation between pixels from two images. Correlation between pixels from two fluorescent recordings of an image suggests spatial colocalization of labeled proteins. GxTX-594, and Kv2.1-GFP or Kv2.2-GFP, had PCC that averaged 0.86 or 0.80, respectively, indicating a high degree of colocalization of both Kv2 proteins with GxTX-594 (Fig 3 B).

We also assessed if GxTX-594 has a preference for either of the Kv2 proteins under these labeling conditions. We calculated the ratio of GxTX-594 to GFP fluorescence intensity from GxTX-594 and Kv2.1-GFP or Kv2.2-GFP. The ratio of GxTX-594 to GFP fluorescence intensity was similar for Kv2.1-GFP or Kv2.2-GFP (Fig 3 C), consistent with the lack of discrimination of GxTX between Kv2.1 and Kv2.2 (Herrington, Zhou et al. 2006).

The colocalization of GxTX-594 specifically with Kv2 proteins makes it clear that the dominant GxTX-594 fluorescence signal from CHO cells is due to binding of Kv2 proteins. However, GxTX accesses the membrane-embedded voltage sensors of Kv2 proteins by partitioning into the outer leaflet of the plasma membrane bilayer (Milescu, Bosmans et al. 2009, Gupta, Zamanian et al. 2015), and a GxTX derivative labeled with an environment sensing fluorophore is detectable in the membrane of this same cell line under similar conditions (Fletcher-Taylor, Thapa et al. 2020). We wondered if GxTX-594 partitioning into CHO cell membranes could contribute to fluorescence measurements conducted throughout this study. To assess GxTX-594 partitioning into the membrane, under conditions used for voltage clamp fluorimetry later in this study, we compared fluorescence from non-transfected CHO cells with cells transfected with Kv2-GFP proteins. The membrane of all cells was identified using wheat germ agglutinin (WGA-405) as a mask to manually draw a region of interest (ROI) where the subsequent analysis was performed. We saw no indication of GxTX-594 enrichment at the plasma membrane of non-transfected cells (Fig 4 A). GxTX-594 fluorescence from non-transfected cells had lower intensities than the extracellular solution (Fig 4 B). We expect lower fluorescence inside of cells as GxTX-594 is membrane impermeant and excludes the 9 nM GxTX-594 in the extracellular solution from the cell interior. When GxTX-594 fluorescence was plotted against Kv2.1-GFP fluorescence, a linear trend was seen with increasing Kv2.1-GFP and GxTX-594 signal (Fig 4 B). Thus GxTX-594 fluorescence intensity from CHO cells is dependent on the presence of Kv2.1 proteins and not membrane partitioning. These results indicate that GxTX-594 partitioning into CHO cell membranes contributes little to the fluorescence measurements made throughout this manuscript.

**Figure 4:**
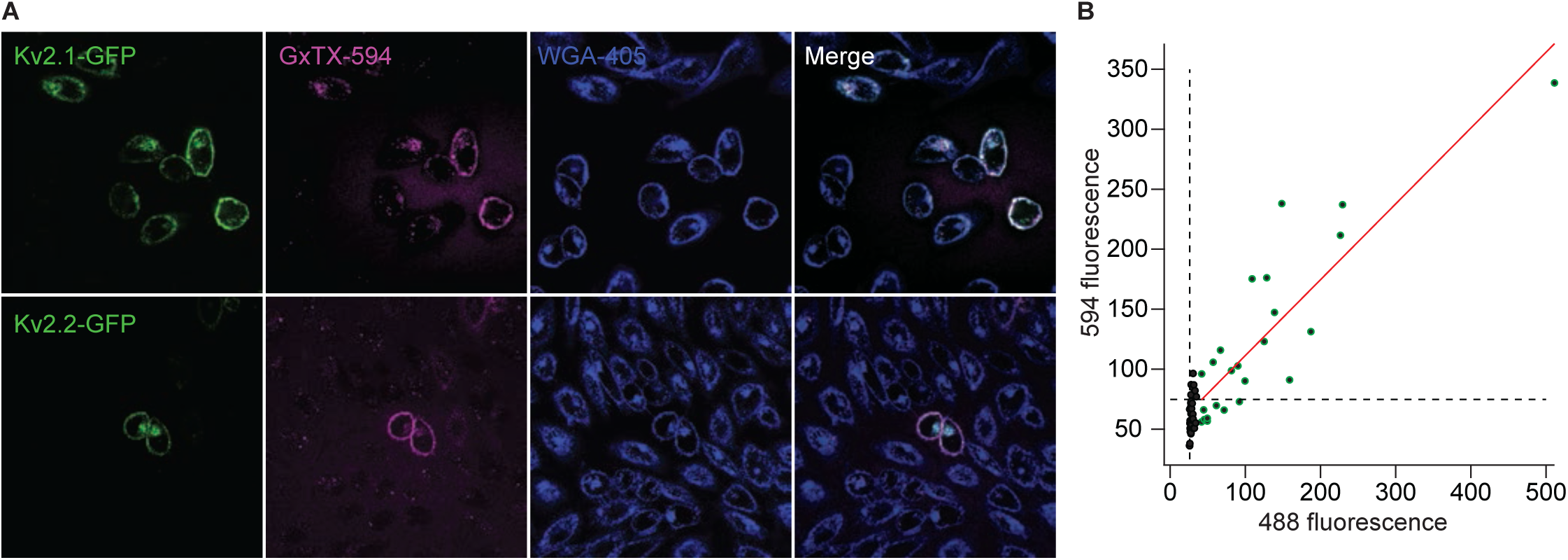
GxTX-594 labeling of surface membranes requires Kv2 protein. A. Fluorescence from live CHO cells transfected with Kv2.1-GFP (upper row) or Kv2.2-GFP (lower row) and labeled with GxTX-594. Airy disk imaging was at a plane above the glass-adhered surface. Cells were incubated with 100 nM GxTX-594 and 5 μg/mL WGA-405 then diluted to 9 nM GxTX-594 and 0.45 μg/mL WGA-405 before imaging. Scale bars are 20 µm. Fluorescence shown was excited by a 594 nm laser (Column 1), 488 nm laser (Column 2) or 405 nm laser (Column 3). Scale bars are 20 µm. B. Fluorescence intensity from 594 nm excitation versus 488 nm excitation. Cell preparation was transfected with Kv2.1-GFP. Each point represents one cell. Cells with obvious GFP fluorescence are green points, cells without are black points. Mean background fluorescence from a region that did not contain cells is indicated by dashed lines. Red line represents a linear fit of cells with obvious GFP fluorescence.

### GxTX-594 modulates Kv2.1 conductance

We performed electrophysiological analyses to determine if GxTX-594 retains the ability to allosterically modulate Kv2.1. GxTX is a partial inverse agonist of Kv2.1 which lowers channel open probability by stabilizing voltage sensors in a resting conformation. Consequently, more positive intracellular voltage is required to activate voltage sensors and achieve the same open probability as without GxTX (Tilley, Angueyra et al. 2019). Previously we estimated that the Kv2.1 complex with GxTX bound to the voltage sensor is at least 5400-fold more stable when the voltage sensor is in its resting conformation (Tilley, Angueyra et al. 2019). Kv2.1 requires more positive intracellular voltage to attain the same level of conductance when bound by GxTX. To characterize the efficacy of GxTX-594 in allosterically modulating Kv2.1 gating, we measured whole cell Kv2.1 ionic currents by voltage clamp in the presence and absence of GxTX-594. From these currents we analyzed the Kv2.1 conductance-voltage (GV) relation by fitting with a 4^th^ power Boltzmann function. The voltage at which the conductance of the fitted function is 50% of maximum, V_mid_, was +73 mV for 100 nM GxTX-594 (Fig 5). For comparison, the V_mid_ of 100 nM GxTX was +67 mV (Tilley, Angueyra et al. 2019). This small difference in V_mid_ between GxTX and GxTX-594 could potentially result from the different extracellular solutions (see methods), differences in allostery or affinities between GxTX and GxTX-594, or the inherent variability of Kv2.1 GVs between cells. The efficacy of GxTX-594 in shifting the GV indicates that it retains efficacy against Kv2.1 and suggests GxTX-594 acts by the same mechanism as GxTX.

**Figure 5:**
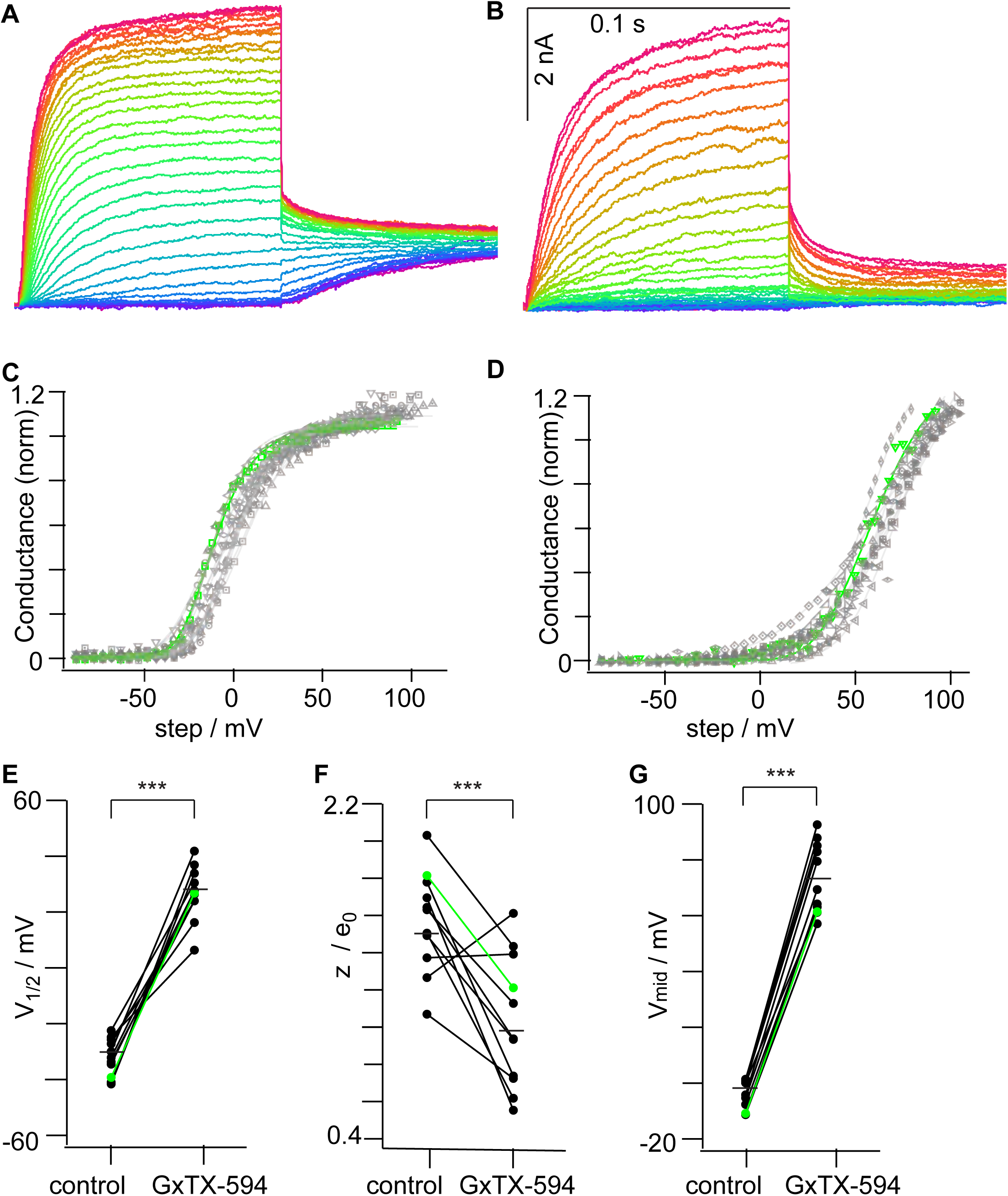
GxTX-594 modulates Kv2.1 conductance. A. Representative Kv2.1-CHO current response under whole cell voltage-clamp. Cells were given 100 ms, 5 mV increment voltage steps ranging from −80 mV (blue) to +120 mV (red) and then stepped to 0 mV to record tail currents. The holding potential was −100 mV. B. Kv2.1 currents from the same cell 5 minutes after the addition of 100 nM GxTX-594. C. Normalized conductance–voltage relationships from Kv2.1 current before application of GxTX-594 (n = 13). Different symbols correspond to individual cells and the green corresponds to cell in panel A. D. Normalized conductance–voltage relationships in 100 nM GxTX-594 (n = 11). E. Mean midpoint of each of 4 independent voltage sensors in the 4^th^ power Boltzmann fit (*V_1/2_*) before (−31 ± 6 mV SD) and after (27 ± 10 mV SD) 100 nM GxTX-594. p < 0.0001, two-tailed T-test. F. Mean elementary charge associated with Boltzmann fit (*z*) in units of elementary charge before (1.5 ± 0.3 e^+^ SD) and after (1.0 ± 0.4 e^+^ SD) 100 nM GxTX-594. p = 0.0013, two-tailed T-test. G. Mean midpoint of conductance change in the 4^th^ power Boltzmann fit (*V_mid_*) before (−2 ± 6 mV SD) and after (73 ± 13 mV SD) 100 nM GxTX-594. p < 0.0001, two-tailed T-test.

### The relationship between GxTX-594 cell surface fluorescence and Kv2.1 voltage activation

To study the relationship between voltage sensor conformational change and GxTX-594 fluorescence, we collected time-lapse fluorescence images from voltage-clamped Kv2.1-CHO cells. We analyzed fluorescence dynamics from airy disk confocal images of cells voltage-clamped in the whole-cell patch clamp configuration. For imaging experiments, Kv2.1-CHO cells were induced for 2 days to express higher densities of Kv2.1 than in Figure 5 ionic current recordings. To minimize voltage clamp error associated with the high level of whole cell Kv2.1 current, the blocker Cs^+^ replaced K^+^ in the pipette solution. As the dose-response experiments indicate GxTX-594 signal equilibrates within 5 min after application of 100 nM GxTX-594 (Fig 2), we adopted a protocol of applying 100 nM GxTX-594 to Kv2.1-CHO cells for 5 minutes and then, to reduce the fluorescence in the bath solution, GxTX-594 was diluted to 9 nM before imaging. The confocal imaging plane containing the most GxTX-594 fluorescence was the glass-adhered surface of CHO cells. At this imaging plane we conducted time lapse imaging of voltage-clamped CHO cells and observed a decrease in GxTX-594 fluorescence when the membrane voltage was stepped from −80 mV to +40 mV. Upon returning the membrane voltage to −80 mV, GxTX-594 fluorescence increased (Fig 6 A and C). During time lapse imaging of voltage-clamped cells, we noticed that GxTX-594 fluorescence in the center of the glass-adhered surface responded more slowly to voltage changes than the periphery. At −80 mV, at the center of the glass-adhered cell surface, relabeling was incomplete after 500 seconds, however, at the cell periphery relabeling neared completion within 200 seconds (Fig 6 A). To quantify this observation, concentric circular ROIs were drawn with the smallest ROI in the center of the glass-adhered surface. The average fluorescence intensities from each of these ROIs were compared to each other and an ROI at the cell periphery (Fig 6 B and C). We quantified the rate of fluorescence change (*k*_Δ*F*_) by fitting the average fluorescence intensities from each ROI with a monoexponential function:

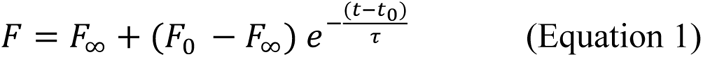

Where τ = 1/*k*_Δ*F*_, *t* = time, *t_0_* = time at start of fit, *F* = fluorescence intensity, *F_0_* = fluorescence at start of fit, *F_∞_* = fluorescence after infinite time. In response to voltage change, the *k*_Δ*F*_ was consistently slower at the center of the glass-adhered surface and faster towards the outer periphery (Fig 6 D and E). While *k*_Δ*F*_ was consistently slower at the center of the glass-adhered surface we observed that the rate of fluorescence change was more pronounced during labeling at −80 mV than unlabeling at 40 mV. When cells were held at a membrane potential of −80 mV, *k*_Δ*F*_ at the periphery was approximately 10-fold faster than the *k*_Δ*F*_ at the center of the cell. In comparison, when the membrane potential was held at 40 mV, *k*_Δ*F*_ at the periphery was approximately 3-fold faster than the *k*_Δ*F*_ at the center of the cell. This difference in kinetics suggests that the concentration of freely diffusing GxTX-594 in the restricted space between the cell membrane and the glass surface is transiently non-uniform after voltage change. Potential reasons for this phenomenon are considered further in the *Limitations of GxTX-594* section of the discussion. We found that the average *k*_Δ*F*_ at −80 mV at the center of the glass-adhered surface was comparable to the *k*_Δ*F*_ at 10 nM from the confluent cells in our dose response experiments (Fig 2 D) at 0.0014 s^-1^ and 0.0011 s^-1^ respectively. This suggests that the Kv2 voltage sensors in the unpatched cells for dose response experiments are the same early resting conformation as voltage-clamped cells at −80 mV (Scholle, Dugarmaa et al. 2004, Tilley, Angueyra et al. 2019). The gradual change in fluorescence intensity over many seconds after a voltage step is inconsistent with a fast electrochromic effect leading to fluorescence change, as the change in fluorescence intensity does not occur instantaneously when the membrane potential is stepped. Additionally, the slowing of *k*_Δ*F*_ in subcellular regions further from the periphery of cells is inconsistent with a slower electrochromic effect, as whole cell voltage clamp should render the membrane surface isopotential.

**Figure 6:**
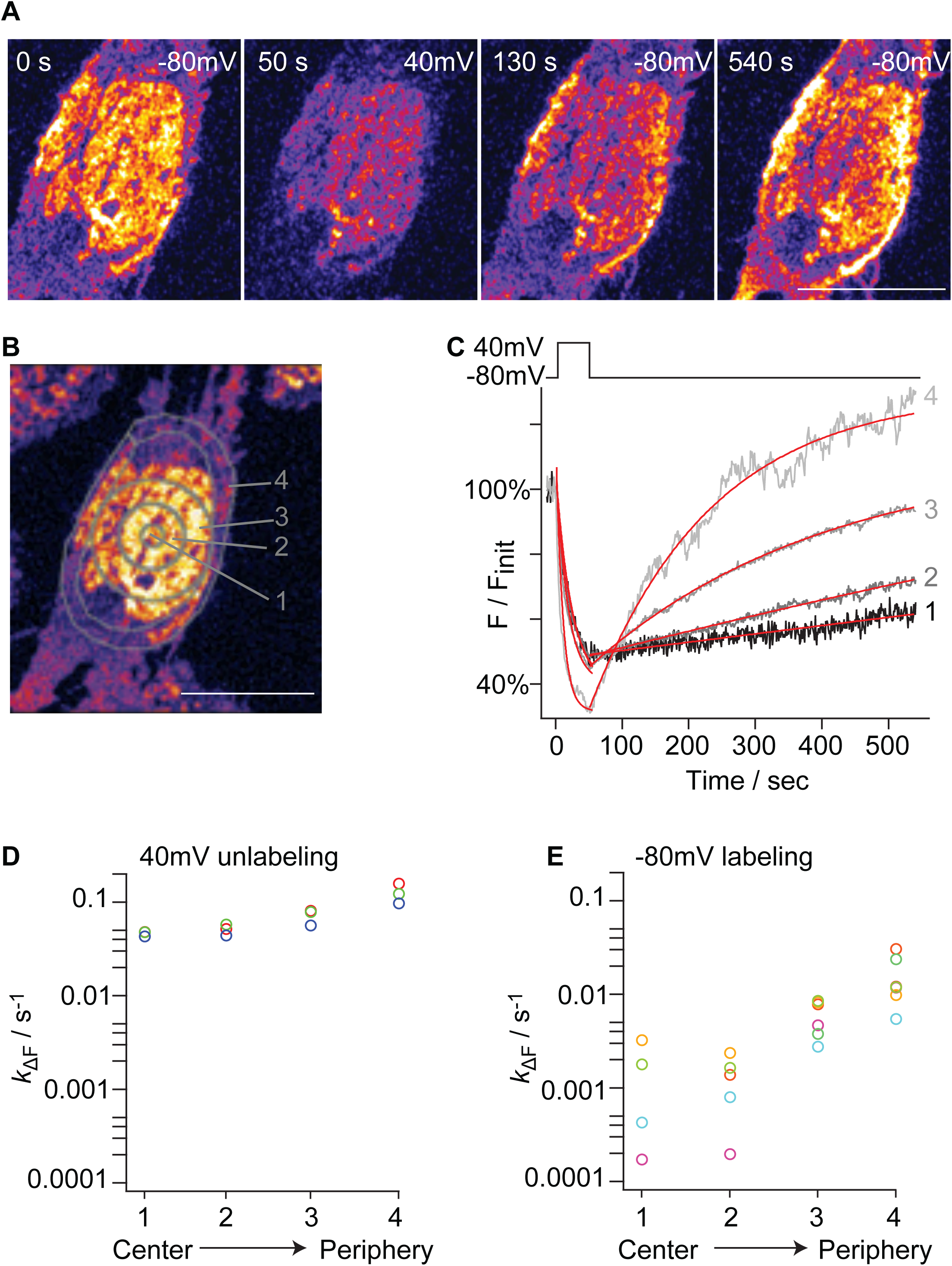
Extracellular access can impact GxTX-594 labeling kinetics. A. Time lapse airy disk images of the glass-adhered surface of a voltage-clamped Kv2.1-CHO cell in 9 nM GxTX-594. Pseudocolored images indicate maximal intensities in white. Time index is in the upper left of each panel and membrane potential is indicated in the upper right. Scale bar in lower right of last panel is 10 µm B. Airy disk image of the glass adhered surface of a voltage-clamped Kv2.1-CHO cell in 9 nM GxTX-594. Gray lines indicate boundaries of ROIs. ROI 1, 2, and 3 are concentric circles each with a respective diameter of 1.8, 4.9, and 9.1 µm. ROI 4 was hand-drawn to contain the apparent cell surface. In all cells analyzed, ROI 1-3 were concentric circles of the same sizes, while ROI 4 varied based on cell shape. Scale bar is 10 µm. C. Representative traces of 594 fluorescence intensity response to voltage changes. Red lines are monoexponential fits (Eq. 1): 40 mV step ROI 1 *k*_Δ*F*_ = 4.29 x 10^-2^ ± 0.26 x 10^-2^ s^-1^, ROI 2 *k*_Δ*F*_ = 4.39 x 10^-2^ ± 0.16 x 10^-2^ s^-1^, ROI 3 *k*_Δ*F*_ = 5.65 x 10^-2^ ± 0.1.5 x 10^-2^ s^-1^, ROI 4 *k*_Δ*F*_ = 9.69 x 10^-2^ ± 0.33 x 10^-2^ s^-1^. −80 mV step ROI 1 *k*_Δ*F*_ = 4.27 x 10^-4^ ± 0.11 x 10^-4^ s^-1^, ROI 2 *k*_Δ*F*_ = 7.999 x 10^-4^ ± 0.053 x 10^-4^ s^-1^, ROI 3 *k*_Δ*F*_ = 2.7556 x 10^-3^ ± 0.0074 x 10^-3^ s^-1^, and ROI 4 *k*_Δ*F*_ = 5.46 x 10^-3^ ± 0.13 x 10^-3^ s^-1^. Background for subtraction was the average intensity of a region that did not contain cells over the time course of the voltage protocol. Each trace was normalized to initial fluorescence intensity before the application of the voltage stimulus. D. Circles indicate *k*_Δ*F*_ at +40 mV from individual cells. Circle coloring indicates data from the same cell. E. Circles indicate *k*_Δ*F*_ at −80 mV from individual cells.

To avoid complications resulting from the spatial variability of kinetics at the glass-adhered surface, further voltage-dependent labeling experiments were conducted while imaging at a plane above the coverslip, where the cell surface has unrestricted access to the bulk extracellular solution. For consistency, we retained the protocol of initially labeling Kv2 channels with 100 nM GxTX-594 and diluting the solution to 9 nM. In retrospect it is not clear that the 100 nM step offers any advantage when imaging at a plane above the glass-adhered surface. Voltage stimuli were given only after fluorescence intensity stabilized at −80 mV; this was at least 9 minutes after dilution of GxTX-594 to 9 nM. We determined the time required for GxTX-594 to reach a stable fluorescence after dilution from 100 nM to 9 nM by time-lapse imaging during dilution (Fig Supplement 2 A, B). The *k*_Δ*F*_ indicated that, on average, equilibration was 85% complete 9 minutes after dilution to 9 nM. After dilution to 9 nM the mean fluorescence intensity decreased by 39.1 ± 8.4% and remained stable (Fig Supplement 2 B).

To investigate the relationship between voltage sensor conformational change and GxTX-594 fluorescence, we measured the fluorescence response of GxTX-594 when the membrane voltage of cells prepared and imaged as described above was stepped from −80 mV to more positive voltages that ranged from −40 mV to +80 mV (Fig 7 A). ROIs corresponding to the cell surface were manually identified and average fluorescence intensity quantified from time-lapse sequences. Both the amplitude and kinetics of fluorescence change from cell surface ROIs are sensitive to voltage (Fig 7 B), similar to prior findings with GxTX-550 (Tilley, Eum et al. 2014). The dynamic fluorescence responses of GxTX-594 to voltage changes occurred at physiologically relevant potentials, suggesting that changes in GxTX-594 labeling intensity or rate could occur in response to changes in cellular electrical signaling. We observed that the voltage-dependent reduction in fluorescence equilibrates at a value above background. Even when cells were given a +80 mV depolarizing voltage stimulus, some GxTX-594 fluorescence remained (Fig 7 B). Most of the residual fluorescence appeared to be localized to the cell surface membrane (Fig 7 A), and the amount varied between cells (Fig 7 C). As such surface labeling was not present on CHO cells in the absence of Kv2 proteins (Fig 4), we consider this residual fluorescence to originate from voltage-insensitive Kv2 proteins. These voltage-insensitive Kv2 proteins could potentially be at the cell surface and have immobilized voltage sensor which are stuck in a resting conformation, or they could be internalized Kv2.1–GxTX-594 complexes that remain just under the cell surface. Kv2.1 channels in HEK293 cells are internalized within 10 minutes at 37^°^C (Deutsch, Weigel et al. 2012, Weigel, Fox et al. 2012, Fox, Haberkorn et al. 2013, Weigel, Tamkun et al. 2013).

**Figure 7:**
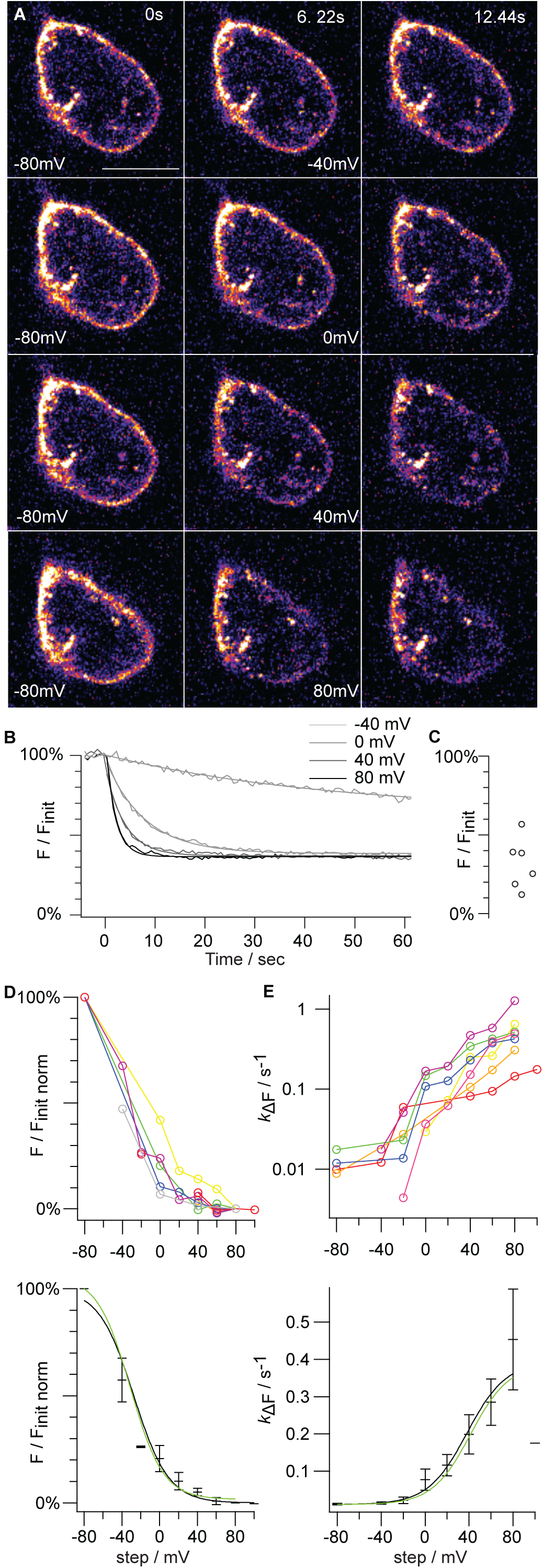
GxTX-594 labeling responds to transmembrane voltage. A. Fluorescence intensities from an optical section of a voltage-clamped Kv2.1-CHO cell in 9 nM GxTX-594. Pseudocolored images indicate maximal intensities in white. Rows indicate voltage step taken from a holding potential of −80 mV. Columns indicate time after voltage step. Scale bar is 10 µm. B. GxTX-594 fluorescence during steps to indicated voltages. Smooth lines are monoexponential fits (Eq. 1): −40 mV *k*_Δ*F*_ = 2.15 x 10^-2^ ± 0.22 x 10^-2^ s^-1^: 0 mV *k*_Δ*F*_ = 1.279 x 10^-1^ ± 0.023 x 10^-1^ s^-1^, 40 mV *k*_Δ*F*_ = 2.492 x 10^-1^ ± 0.062 x 10^-1^ s^-1^ and 80 mV *k*_Δ*F*_= 4.20 x 10^-1^ ± 0.11 x 10^-1^ s^-1^. ROIs were hand-drawn around the apparent cell surface membrane based on GxTX-594 fluorescence. 0% was set by subtraction of background which was the average intensity of a region that did not contain cells over the time course of the voltage protocol. For each trace, 100% was set from the initial fluorescence intensity at −80 mV before the subsequent voltage step. C. Fluorescence intensity remaining at the end of 50 s steps to +80 mV. Each circle represents one cell. Background subtraction as in panel B. D. Voltage-dependence of fluorescence intensity at the end of 50 s steps. For each cell, 100% was set from the initial fluorescence intensity at −80 mV before the first step to another voltage. Cells did not always recover to initial fluorescence intensity during the −80 mV holding period between voltage steps. (Top panel) Circle coloring indicates data from the same cell and lines connect points from the same cell. Gray circles represent data shown in B. (Bottom panel) black bars represent the mean *F*/*F_init_* at each voltage and error bars represent the standard error of the mean. Black line is fit of a first-order Boltzmann equation (Eq. 2): *V_1/2_* = −27.4 ± 2.5 mV, *z* = 1.38 ± 0.13 *e*_0_. Green line is prediction from the EVAP model at 9 nM GxTX. E. Voltage dependence of fluorescence intensity kinetics (*k*_Δ*F*_). (Top panel) circle coloring is the same as panel D. (Bottom panel) black bars represent the average *k*_Δ*F*_ at each voltage and error bars represent the standard error of the mean. Black line is a first-order Boltzmann equation fit to the *k*_Δ*F*_–voltage relation: *V_1/2_* = +38 ± 15 mV, *z* = 1.43 ± 0.35 *e*_0_. Green line is prediction from the EVAP model at 9 nM GxTX. Error bars are standard error of the mean.

To compare voltage-response properties between cells, we used a normalization procedure to analyze only the voltage-sensitive fraction of the fluorescence from each cell, which we defined as the fluorescence that changed between −80 mV and +80 mV. The initial fluorescence at a holding potential of −80 mV was normalized to 100% *F/F_init_*, and residual fluorescence after a +80 mV step was normalized to 0% *F/F_init_* (Fig 7 D). To characterize the voltage dependence of the Kv2.1–GxTX-594 interaction, fluorescence–voltage (*FV*) responses were fit with a

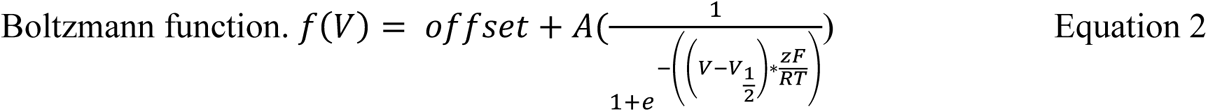

Where *V* = voltage, *offset* = the offset from zero of the Boltzmann distribution, *A* = the amplitude, *z* = number of elementary charges, *F* = Faraday’s constant, *R* = the universal gas constant, *T* = temperature (held at 295 K). This fit had a half maximal voltage midpoint (*V_1/2_*) of −27 mV and a steepness (*z*) of 1.4 elementary charges (*e*_0_) (Fig 7 D lower panel, black line). This is similar to a fit of voltage sensor movement in Kv2.1-CHO cells, without any GxTX present: *V_1/2_* = −26 mV, *z* = 1.6 *e*_0_ (Tilley, Angueyra et al. 2019).

To determine the temporal response of GxTX-594 labeling to voltage change, we compared rates of fluorescence change (*k*_Δ*F*_) at varying step potentials (Fig 7 E) in the same data set that were analyzed in Figure 7 D. We quantified *k*_Δ*F*_ by fitting the average fluorescence from voltage-clamped cells with a monoexponential function (Eq 1).

In response to voltage steps from a holding potential of −80 mV to more positive potentials, *k*_Δ*F*_ increased progressively as step potential was increased. After a return to −80 mV, labeling occurred at a rate similar to unlabeling at −40 mV, indicating that the rate of relaxation to a labeled/unlabeled equilibrium saturates below −40 mV. We noted the degree of variability in *k*_Δ*F*_ measurements became greater at more positive potentials (Fig 7 E top and bottom panels). At −80 mV there was a 2-fold range in *k*_Δ*F*_ values and a 9-fold range at +80 mV. The relatively low variation in *k*_Δ*F*_ at −80 mV suggests that fits of the upward relaxation at −80 mV are relatively consistent in this data set, despite variance in fluorescence intensity after rebinding (Fig 6 C and Fig 9B). Possible reasons for cell-to-cell variability at more positive voltages are discussed further in the *Limitations of GxTX-594* section of the Discussion. The average *k*_Δ*F*_–voltage response was fit with a Boltzmann distribution (Eq. 2) resulting in *V_1/2_* = +38 ± 15 mV and *z* = 1.43 ± 0.35 *e*_0_ (Fig 7 E, black line lower panel). While the precision of this fit is limited by the variability in *k*_Δ*F*_ values, it is notable that the *V_1/2_* and *z* from the Boltzmann fit is similar to the voltage dependence of integrated gating charge of the GxTX–Kv2.1 complex: *V_1/2_* = +41.3 mV, *z* = 1.5 *e*_0_ (Tilley, Angueyra et al. 2019). We later propose a mechanistic underpinning for this correlation.

This analysis suggests that the dynamics of GxTX-594 labeling convey a surprisingly straightforward readout of Kv2.1 voltage sensor activation: the magnitude of the response is determined by the probability of voltage sensor activation of Kv2.1 alone and the kinetics of the response are determined by the probability of voltage sensor activation in the Kv2.1–GxTX-594 complex.

### Temperature dependence of fluorescence kinetics

As part of our efforts to identify the underpinnings of the substantial variation in GxTX-594 *k*_Δ*F*_ measurements, we assessed the temperature dependence of *k*_Δ*F*_. Given the marked temperature sensitivity reported for Kv2.1 conductance (Yang and Zheng 2014), we wondered if this could be due to the fluctuations in ambient temperature (27-29°C). To assess the temperature dependence of GxTX-594 labeling, the cell bath solution was held at either 27°C or 37°C and stepped to 0 mV for a measurement of *k*_Δ*F*_. The average change in *k*_Δ*F*_ over this 10 °C difference, or Q_10_, was 3.8-fold (Fig Supplement 3). This Q_10_ predicts only a 1.3-fold increase in *k*_Δ*F*_ can be attributed to a 2°C variation, suggesting only a small fraction of the 6-fold variability could be attributed to temperature fluctuations. Other potential sources of variability are considered in the *Limitations of GxTX-594* section of the discussion.

### The relation between voltage sensor activation and GxTX-594 dynamics can be recapitulated by rate theory modeling

To enable translation of the intensity of fluorescence from GxTX-594 on a cell surface into a measure of conformational change, we developed an EVAP model, a series of equations derived from rate theory that relate cell labeling to voltage sensor activation. The framework of the EVAP model is generalizable to fluorescent molecular probes that report conformational changes by a change in binding affinity. In the EVAP model, the proportion of labeled vs. unlabeled Kv2 in a membrane is determined by the proportion of voltage sensors in resting vs. activated conformations. The model assumes the innate voltage sensitivity of the Kv2 subunit is solely responsible for controlling voltage dependence. EVAP labeling is voltage-dependent because the binding and unbinding rates are different for resting and activated conformations of voltage sensors.

Voltage-activation of Kv2 channels involves many conformational changes (Scholle, Dugarmaa et al. 2004, Jara-Oseguera, Ishida et al. 2011, Tilley, Angueyra et al. 2019). However, models which presume independent activation of a voltage sensor in each of the four Kv2.1 subunits accurately predict many aspects of voltage activation and voltage sensor toxin binding (Lee, Wang et al. 2003, Tilley, Angueyra et al. 2019). For simplicity, we model Kv2 proteins as having only resting and activated conformations. With the simplifying assumptions that the conformational change in each voltage sensor is independent, a rate theory model was developed consisting of 4 interconnected states (Scheme I: EVAP Allosteric Expansion).

**Scheme I:**
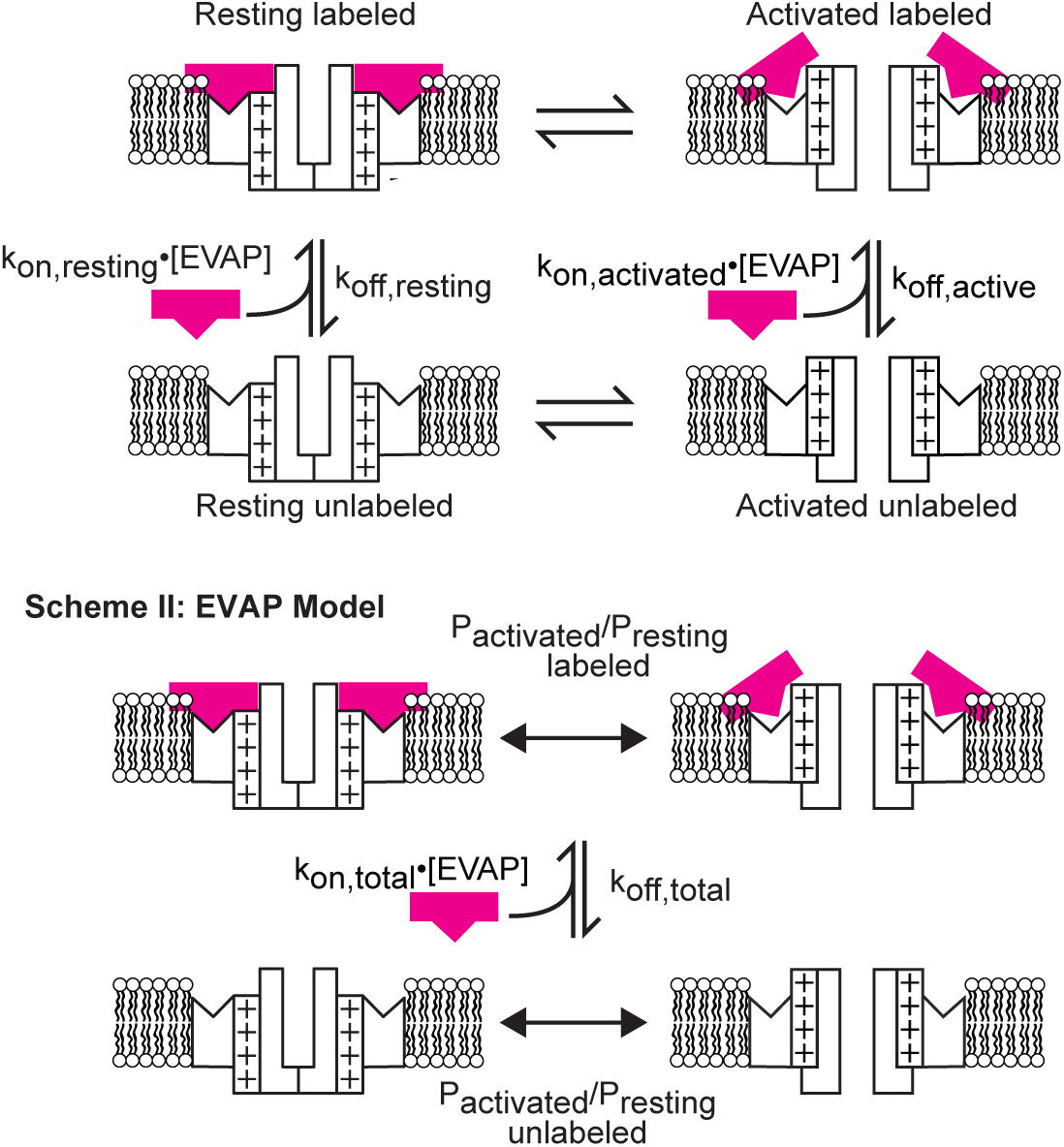
EVAP Allosteric Expansion.

When voltage sensors change from resting to activated conformations, the binding rate of the GxTX-594 EVAP decreases and the unbinding rate increases. When the membrane voltage is held constant for sufficient time, the proportions of labeled and unlabeled proteins reach an equilibrium. EVAP labeling requires seconds to equilibrate (Fig 7), whereas Kv2 channel gating equilibrates in milliseconds (Tilley, Angueyra et al. 2019), three orders of magnitude more quickly. These distinct time scales of equilibration suggest an approximation to model the EVAP labeling response: voltage sensor conformations achieve equilibrium quickly such that only their distribution at equilibrium is expected to greatly impact the kinetics of labeling and unlabeling, allowing Scheme I to collapse into Scheme II, which depicts the structure of the EVAP Model used for calculations.

We generated a model of dynamic GxTX-594 fluorescence using the EVAP model and empirical measurements. In the EVAP model, at any given voltage, there is a probability that a voltage sensor is either in its resting conformation (*P_resting_*), or in its activated conformation (*P_activated_*), such that *P_activated_* = (1 – *P_resting_*). The equilibrium for voltage sensor activation is then a ratio of activated to resting voltage sensors (*P_activated_*/*P_resting_*) in which

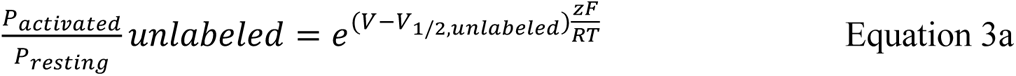

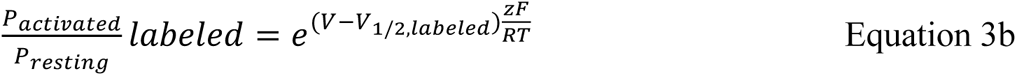

where *V_1/2_* is the voltage where *P_activated_*/*P_resting,_* = 1. In a prior study, our analysis of the conductance–voltage relation of Kv2.1 yielded a *V_1/2_* = −32 mV with *z* = 1.5 *e*_0_ for the early movement of 4 independent voltage sensors, and found that with a saturating concentration of GxTX the *V_1/2_* = +42 mV (Tilley, Angueyra *et al*. 2019, Fig. 1 C). These values were used for *V_1/2,unlabeled_, z, and V_1/2,labeled_* respectively (Table 1). To relate voltage sensor activation to labeling and unlabeling, we used microscopic binding (*k_on_*[*EVAP*]) and unbinding (*k_off_*) rates that are distinct for resting and activated voltage sensors. We estimated values for these rates assuming

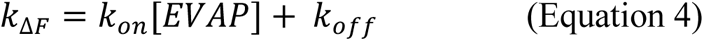

**Table 1.** Parameters used for calculations to generate Scheme I: EVAP Allosteric Expansion.

| Parameter | Value |
| --- | --- |
| $k_{on,resting}$ | $0.30 \mu\text{M}^{-1} \text{s}^{-1}$ |
| $k_{off,resting}$ | $0.0081 \text{s}^{-1}$ |
| $k_{on,activated}$ | $0.21 \mu\text{M}^{-1} \text{s}^{-1}$ |
| $k_{off,activated}$ | $0.39 \text{s}^{-1}$ |
| $V_{1/2,unlabeled}$ | $-32 \text{ mV}$ |
| $V_{1/2,labeled}$ | $41 \text{ mV}$ |
| $z$ | $1.5 e_0$ |

and

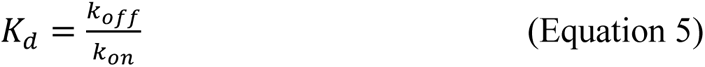

To calculate the *k_on,resting_* and *k_off,resting_* values reported in Table 1, we used the saturating value at negative voltages of the *k*_Δ*F*_–voltage relation (Fig 7 E), and *K_d_* from dose-response imaging (Fig 2 D). In 9 nM GxTX-594, at greater than +40 mV, voltage-dependent unlabeling was nearly complete, indicating that *k_off,activated_* >> *k_on,activated_*[*EVAP*]. We input the saturation of *k*_Δ*F*_ at positive voltages as *k_off,activated_* (Fig 7 E). The slow labeling of activated voltage sensors confounded attempts to measure *k_on,activated_* directly, and we used the statistical thermodynamic principle of microscopic reversibility (Lewis 1925) to constrain *k_on,activated_*:

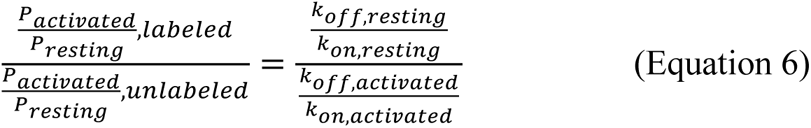

The EVAP model (Scheme II) has only a single microscopic binding rate, *k_on,total_*[*EVAP*], and unbinding rate, *k_off,total_*. *k_on,total_* is a weighted sum of both *k_on,resting_* and *k_on,activated_* from Scheme I. The weights for *k_on,total_* are the relative probabilities that unlabeled voltage sensors are resting or activated, which is determined at any static voltage by an equilibrium constant, 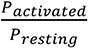 unlabeled:

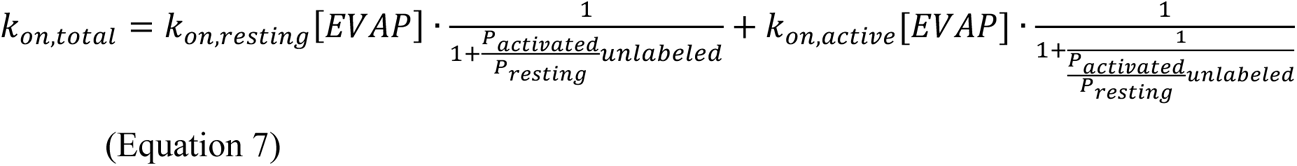

Similarly, *k_off,total_* is determined by the unbinding rate from resting voltage sensors (*k_off,resting_*) and the unbinding rate from activated voltage sensors (*k_off,activated_*), and weighted such that:

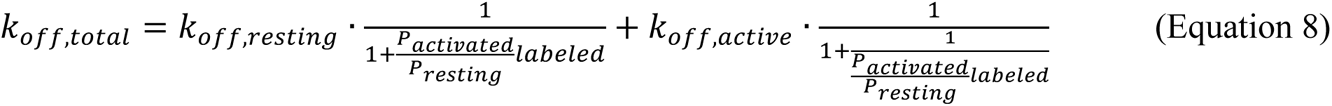

Using *k_on,total_* and *k_off,total_* we can compute *k*_Δ*F*_ using Equation 4. To test the predictive value of the EVAP model, we compared the predicted *k*_Δ*F*_ to empirical measurements in 9 nM GxTX-594 and found the EVAP model (Scheme II) had a *V_1/2_* and *z* that differed by only 3.24 mV and 0.05 *e*_0_ respectively (Fig 7 E, blue line, lower panel).

The EVAP model was also used to predict the magnitude of GxTX-594 fluorescence changes on cell surfaces. In theory, the ratio of fluorescence at a test voltage to fluorescence at a prior voltage (*F/F*_*init*_) is equal to the probability that a Kv2 subunit is labeled by GxTX-594 (*P_labeled_*):

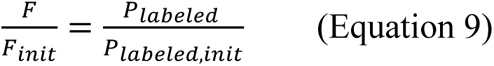

The equilibrium *P_labeled_* at any voltage can be determined from microscopic binding rates associated with Scheme II where:

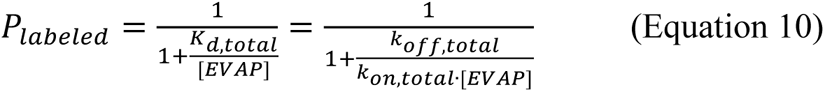

Using this model, we calculated the fluorescence–voltage relation from *P_labeled_* (Equation 9) and compared these calculated values to our experimental data. The model predicts fluorescence response with a *V_1/2_* and *z* that differed by only −4.05 mV and 0.06 *e*_0_ respectively (Fig 7 D, blue line, lower panel).

### The EVAP model predicts that changes in labeling report conformational changes of unlabeled Kv2 proteins

We use the EVAP model to investigate general principles of the relation between voltage sensor activation and labeling. In particular, we are interested in exploiting EVAP labeling to reveal information about voltage sensor activation in the unlabeled Kv2 population, akin to how Ca^2+^ indicators are used to reveal the concentration of free Ca^2+^. The EVAP model was used to predict labeling at a series of EVAP concentrations (Fig 8 A). The model does not include EVAP signal that is insensitive to voltage (Fig 7 C). EVAP labeling is not directly sensitive to voltage but is determined by the probability that voltage sensors are in an activated conformation (Fig 8 A). As GxTX-594 binds resting voltage sensors much more rapidly than activated voltage sensors (*k_on,resting_* >> *k_on,activated_*), the probability that unlabeled voltage sensors are resting determines the labeling rate, and influences *P_bound_*. The rate of fluorescence change (*k*_Δ*F*_) in response to variation in membrane potential is determined by the probability that voltage sensors in the Kv2.1–GxTX-594 complexes are at rest (Fig 8 B). Thus *k*_Δ*F*_ is not readily interpreted as a conformational change in unlabeled Kv2 proteins. Since *k*_Δ*F*_ = *k_on_*[GxTX-594] + *k_off_*, concentrations of GxTX-594 where *k_on_*[GxTX-594] << *k_off_* will cause *k*_Δ*F*_ to be determined by *k_off_* and will have little dependence on the concentration of GxTX-594. The EVAP model predicts that as GxTX-594 concentration decreases, the rate of fluorescence change *k*_Δ*F*_ asymptotically approaches the probability that labelled channels are active (Fig 8 B). However, the EVAP model predicts that as GxTX-594 concentration decreases, the change in labeling (*ΔF*/*ΔF_max_*) asymptotically approaches the probability that unlabeled voltage sensors are resting (Fig 8 A). Even at 100 nM GxTX-594, more than 3 times the *K_d_* for the resting conformation, the midpoint of the Boltzmann distribution describing the change in labeling differs from the midpoint of the QV of unlabeled Kv2.1 proteins by only 25 mV. Thus, the thermodynamics of the ensemble of the free EVAP and labeled and unlabeled Kv2 proteins undergird a remarkable and readily interpretable phenomenon: a decrease in GxTX-594 fluorescence indicates a decrease in the number of unlabeled voltage sensors that are at rest. Thus, a simple qualitative interpretation can be made: a GxTX-594 fluorescence change means unlabeled Kv2 voltage sensors have changed conformation.

**Figure 8:**
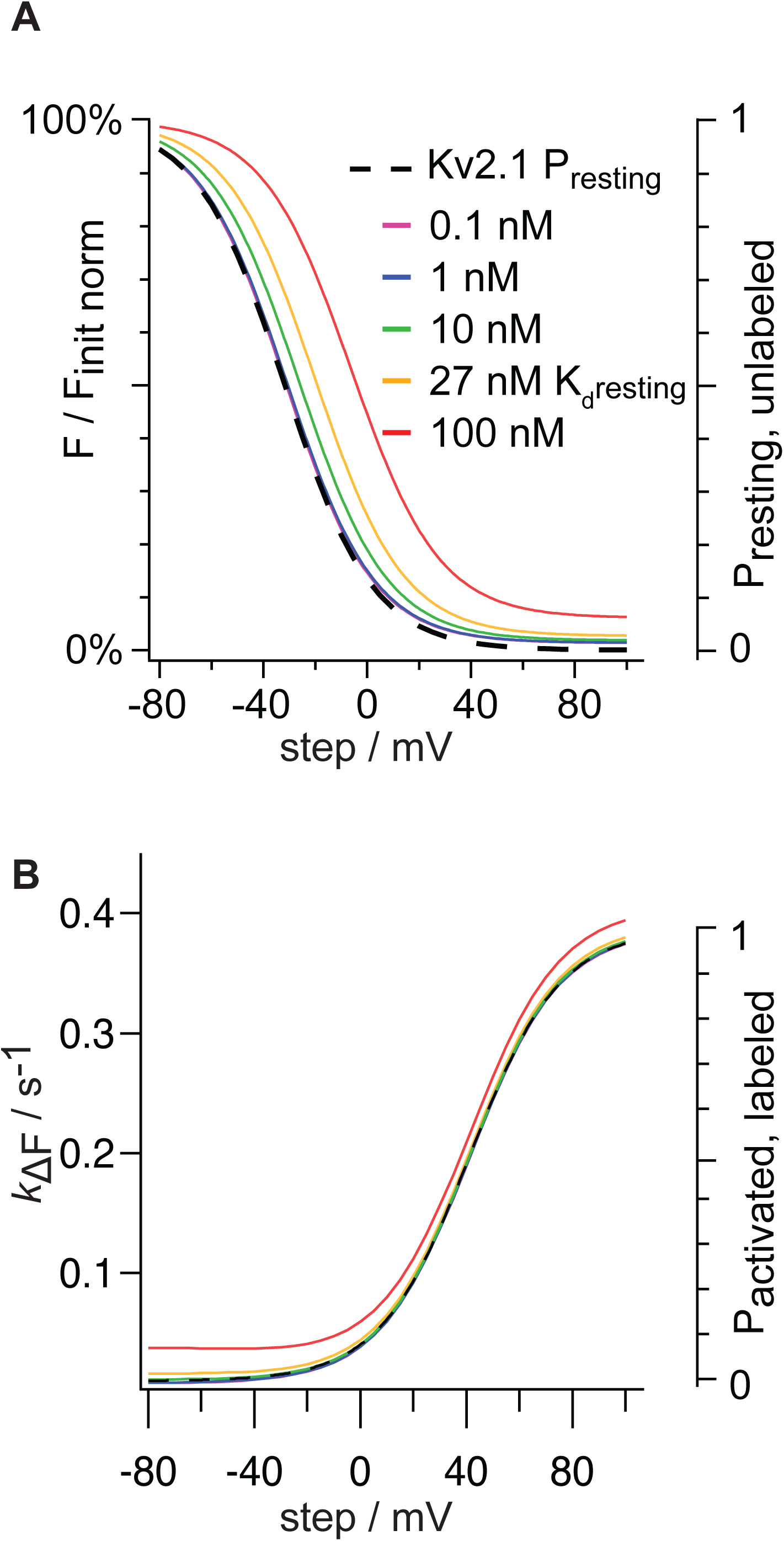
Relationship of GxTX-594 labeling to probability of Kv2 voltage sensor activation. A. EVAP model predictions of voltage-dependence of cell surface fluorescence intensity at different concentrations of EVAP in solution. Bottom axis represents membrane voltage. The left axis represents the predicted fluorescence relative to the situation where all voltage sensors are at rest and does not include EVAP signal that is insensitive to voltage. The right axis represents the probability that voltage sensors of unlabeled Kv2.1 are in their resting conformation. B. EVAP model predictions of concentration- and voltage-dependence of *k*_Δ*F*_. Colors correspond to panel A. The right axis represents the probability that voltage sensors of labeled Kv2.1 are in their active conformation.

### Repetitive action potential-like stimuli amplify the GxTX-594 response

Action potentials involving Kv2 currents occur on the millisecond time scale (Liu and Bean 2014, Kimm, Khaliq et al. 2015), orders of magnitude faster than the GxTX-594 response. Thus, the slow EVAP response will integrate Kv2 conformations occurring over many seconds. The voltage waveforms that are, arguably, most relevant to Kv2 function are repetitive action potentials. Kv2 currents can profoundly impact the firing frequency of neurons and the ability of neurons to sustain high frequency firing (Liu and Bean 2014, Kimm, Khaliq et al. 2015). The particular roles of Kv2 currents in high frequency firing are underpinned by their deactivation kinetics, which are much slower than other Kv currents (Liu and Bean 2014). Kv2.1 OFF gating charge movement reveals that voltage sensor deactivation is even slower than pore closure (Tilley, Angueyra et al. 2019). Thus, sufficiently high action potential frequencies can prevent Kv2 proteins from fully deactivating before a next action potential is triggered, creating a kinetic trap that progressively accumulates activated voltage sensors. This behavior of Kv2 proteins predicts that a voltage stimulus of repetitive action potentials could evoke a more robust fluorescence signal than the EVAP model, which assumes continuous equilibrium of voltage sensors, and thus cannot kinetically trap activated conformations.

We crudely mimicked action potentials with trains of 2 ms voltage steps from −80 to +40 mV, and observed the changes in fluorescence on GxTX-594–labeled Kv2.1-CHO cells (Fig 9 A, B). To assess frequency response, step frequency was varied from 0.02 to 200 Hz. Unlabeling occurred and *k*_Δ*F*_ increased with stimulus frequency (Fig 9 C, D). We compared these fluorescence responses to those predicted by the EVAP model. Predictions of *F/F_init_* and *k*_Δ*F*_ were made from the EVAP model by summing the products of time-averaged probability of being at each voltage (*P_VN_*) and the fluorescence change predicted at that voltage (Δ*F_vn_*)

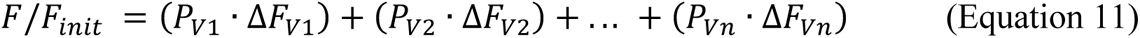

For voltage steps from −80 to +40 mV Eq. 11 is:

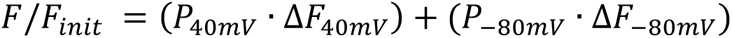

When the stimulus frequency was less than 50 Hz, the EVAP model did a reasonable job of predicting fluorescence change. At and above 50 Hz, fluorescence decreased by more than the EVAP model predicted, even without accounting for a voltage-insensitive fraction of Kv2.1 proteins (Fig 9 C, blue line, lower panel). As discussed above, this divergence from the equilibrium-based EVAP model is expected at frequencies where voltage steps are shorter than Kv2 equilibration times. The time constant of activating gating current decay from Kv2.1-CHO cells was 1.3 ms at +40 mV (Tilley, Angueyra et al. 2019), which means the majority of voltage sensors are effectively activated during the 2 ms +40 mV steps. In contrast, the time constant of deactivating gating current decay from Kv2.1-CHO cells was 22 ms at −80 mV (Tilley, Angueyra et al. 2019), which means Kv2.1 is expected to become kinetically trapped in activated conformations when stimuli to +40 mV from −80 mV are ∼50 Hz or faster. Thus, the amplified EVAP response appears consistent with voltage sensors failing to deactivate before the next stimulus, leading to an accumulation of activated voltage sensors, and a more dramatic fluorescence response than predicted by the EVAP model. An additional factor that could contribute to more EVAP unlabeling than predicted by the EVAP model is the possibility that subsaturating GxTX-594 has little impact on voltage sensor deactivation kinetics. Overall, these dynamics indicate that the magnitude of the change in GxTX-594 fluorescence intensity will be amplified during repetitive action potentials, a regime of electrophysiological signaling where Kv2 currents are critical.

**Figure 9:**
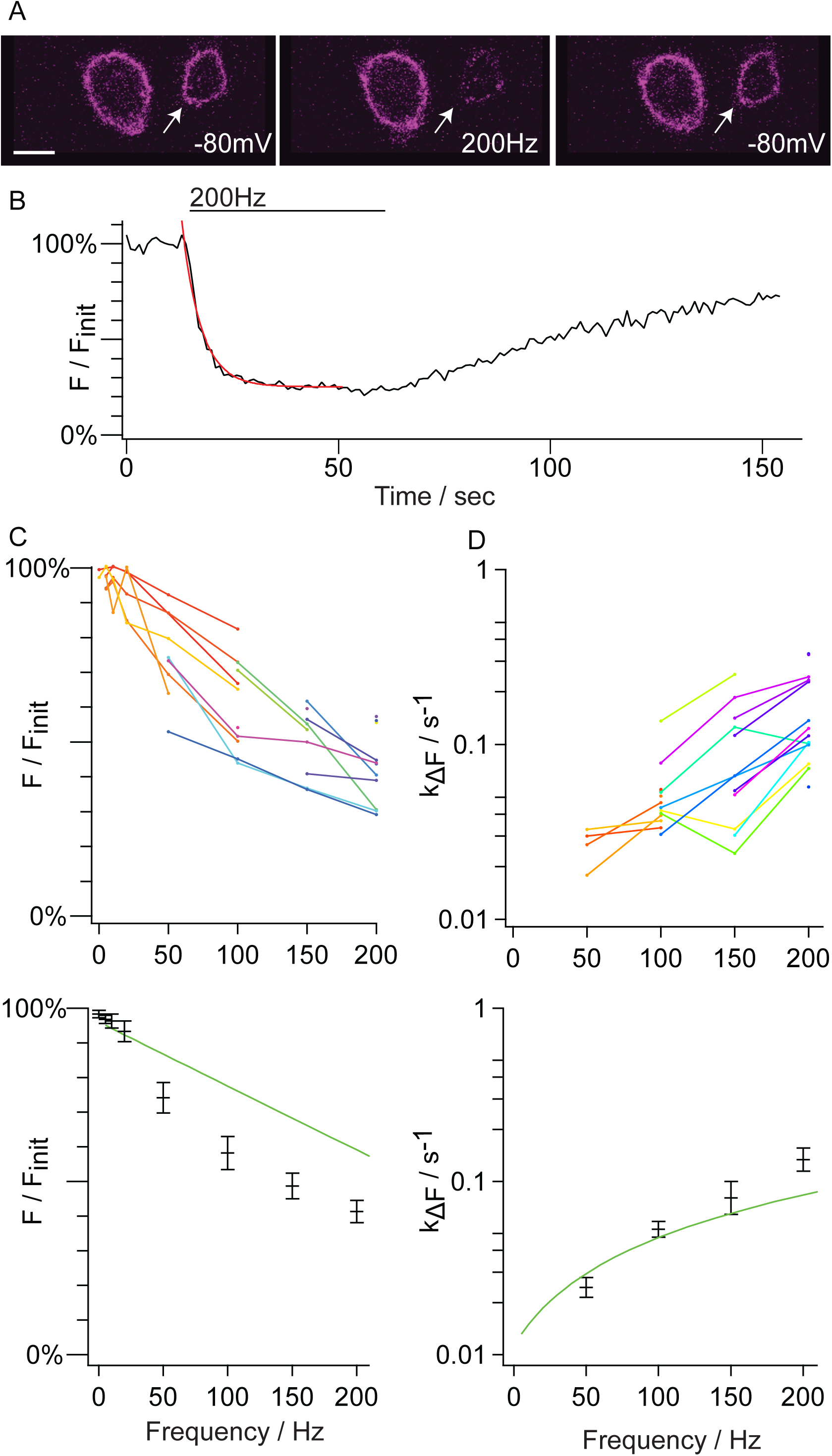
High frequency repetitive stimuli amplify the GxTX-594 response. A. Fluorescence intensity from Kv2.1-CHO cells incubated in 9 nM GxTX-594. Arrow indicates voltage-clamped cell. Fluorescence at holding potential of −80 mV (Left), after 50 s of 200 Hz stimulus (Middle), and 100 s after the cell is returned to holding potential of −80 mV (Right). Stimulus was a 2 ms step to +40 mV. Note that in each panel the unclamped cell (Left cell in each panel) does not show a change in fluorescence. Scale bar is 10 µm. B. Representative trace with GxTX-594 unlabeling at 200 Hz. The trace was fit using a monoexponential (Eq. 1). Fit is overlaid and is in red. The *k*_Δ*F*_ at 200 Hz was obtained from this fit. For 200 Hz *k*_Δ*F*_ = 0.2327 ± 0.0099 x 10^-1^ s^-1^. 0% *F/F_init_* was set by subtraction of the average intensity of a region that did not contain cells. 100% was set by the initial fluorescence intensity at −80 mV. C. Fluorescence–stimulus frequency relation. Points indicate *F/F_init_* from individual cells. (Top panel) point coloring indicates data from the same cell. (Bottom panel) black bars represent the average *F/F_init_* at each voltage and error bars represent the standard error of the mean. Green line is the prediction of the EVAP model at a concentration of 9 nM. D. *k*_Δ*F*_–stimulus frequency relation. Plotted as in panel C.

The EVAP model indicates that GxTX-594 response kinetics are determined by the conformational changes of labeled Kv2 proteins. An assessment of EVAP kinetics in response to increasing frequency of action potential-like voltages, did not reveal major deviations from the EVAP model, in contrast to the fluorescence intensity measurements from the same cells. We predicted EVAP kinetic responses as

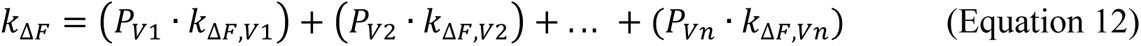

where *P_Vn_* is as in Eq. 11 and *k_ΔF,n_* is *k*_Δ*F*_ at that particular voltage. For voltage steps from −80 to

+40 mV Eq. 12 is:

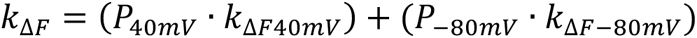

The model produced a frequency dependence of *k*_Δ*F*_ that was within range of the empirical data (Fig 9 D, blue line, lower panel).

This finding is interesting because the response of GxTX-594 labeling to rapidly varying stimuli indicates that the intensity and kinetics of GxTX-594 fluorescence respond distinctly to different voltage waveforms. While the change in fluorescence intensity is expected to be magnified by high-frequency firing, the kinetics of labeling are expected to remain more true to the EVAP model.

### GxTX-594 selectively labels Kv2 proteins

An important consideration for determination of whether GxTX-594 could reveal conformational changes of endogenous Kv2 proteins is whether the EVAP is selective for Kv2 proteins. Electrophysiological studies have concluded that the native GxTX peptide is selective for Kv2 channels, with some off-target modulation of A-type Kv4 channels (Herrington, Zhou et al. 2006, Liu and Bean 2014, Speca, Ogata et al. 2014). However, electrophysiological testing cannot determine whether ligands bind unless they also alter currents (Sack, Stephanopoulos et al. 2013). Furthermore, structural differences between wild-type GxTX and the GxTX-594 variant could potentially alter selectivity among channel protein subtypes.

To test whether GxTX-594 binds other voltage-gated K^+^ channel subtypes, we quantified surface labeling and analyzed colocalization of GxTX-594 with a selection of GFP-labeled voltage-gated K^+^ channel subtypes. A series of GFP-K^+^ channel fusions were identified that express and retain function with a GFP tag: rat Kv4.2-GFP, rat Kv1.5-GFP, and mouse BK-GFP. Transfection of each of these channel subtypes into CHO cells resulted in voltage-dependent outward currents, consistent with cell surface expression of the K^+^ channels (Fig Supplement 1 A). In imaging analysis, the surface membrane was identified by labeling with a fluorescent wheat germ agglutinin (WGA) conjugate. Transfection of all K^+^ channel-GFP constructs resulted in GFP localization at the cell surface, however the majority of the Kv4.2 and Kv1.5 appeared intracellularly retained (Fig Supplement 1 B). Consistent with reports in other cell lines, cell surface localization improved when Kv4.2 was co-transfected with KChIP2 (Shibata, Misonou et al. 2003) or Kv1.5 with Kvβ2 (Shi, Nakahira et al. 1996) (Fig 10 A).

**Figure 10:**
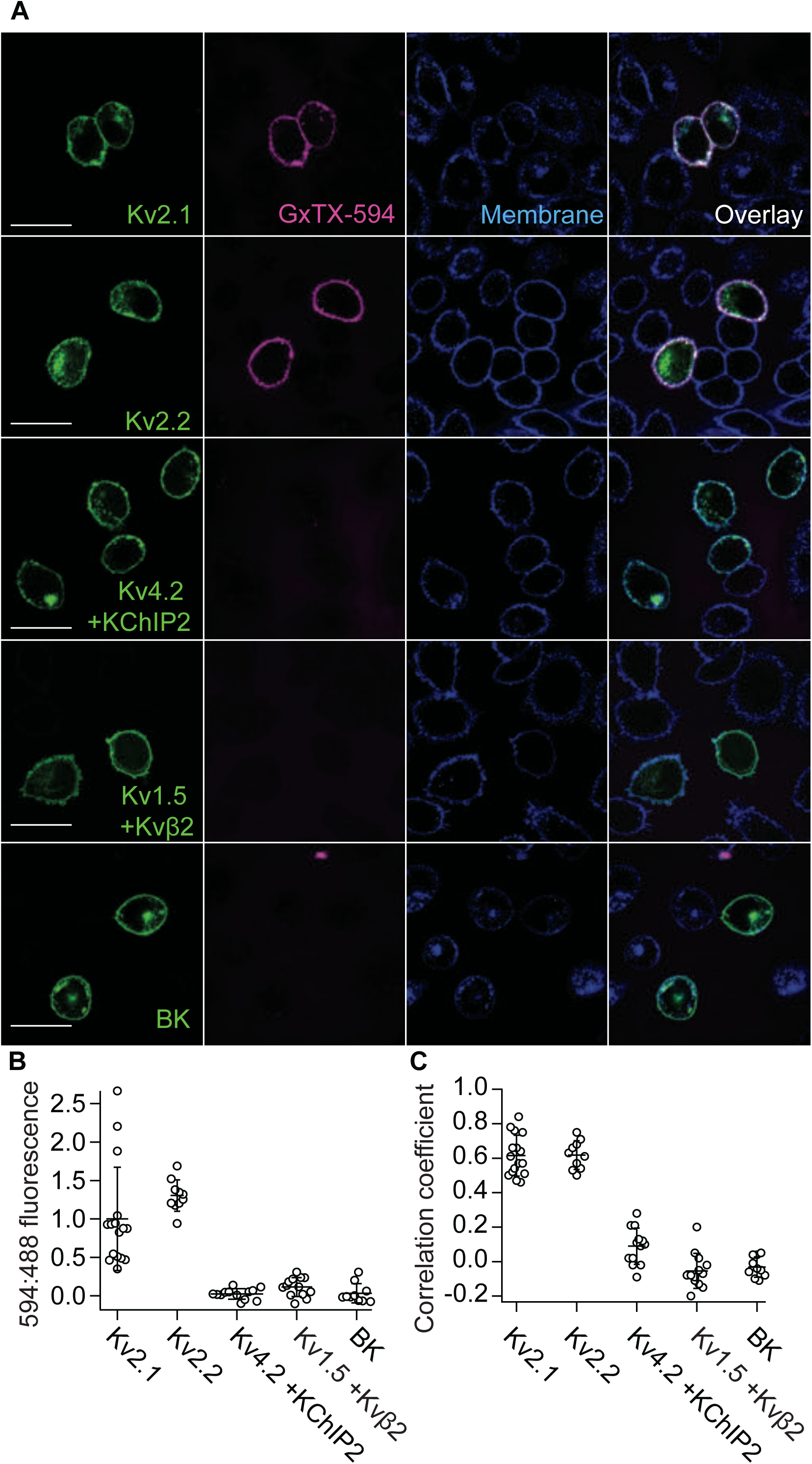
GxTX-594 selectively labels Kv2 proteins. A. Fluorescence from live CHO cells transfected with Kv2.1-GFP, Kv2.2-GFP, Kv4.2-GFP + KChIP2, Kv1.5-GFP + Kvβ2, or BK-GFP (Indicated by row) and labeled with GxTX-594. Airy disk imaging was from a plane above the glass-adhered surface. Cells were incubated with 100 nM GxTX-594 and 5 μg/mL WGA-405 then rinsed before imaging. Fluorescence corresponds to emission of GFP (Column 1), GxTX-594 (Column 2), WGA-405 (Column 3), or an overlay (Column 4). Scale bars are 20 µm. B. Intensity of GxTX-594 labeling for different K^+^ channel-GFP types. Fluorescence intensity resulting from 594 nm excitation of GxTX-594 is divided by fluorescence intensity resulting from 488 nm excitation of GFP. This value was normalized to the average 594:488 ratio from GxTX-594 and Kv2.1-GFP. Circles indicate measurements from individual cells. Only cells with obvious GFP expression were analyzed. For analysis ROIs were drawn around the cell membrane indicated by WGA-405 fluorescence. Pixel intensities were background-subtracted before analyses by subtracting the average fluorescence of a background ROI that did not contain cells from the ROI containing the cell; this occasionally resulted in ROIs with negative intensity. Kv2.1 n = 16, Kv2.2 n = 10, Kv4.2 n = 13, Kv1.5 n = 13, BK n =10; n indicates the number of individual cells analyzed during a single application of GxTX-594. Bars are arithmetic mean. Significant differences were observed between 594:488 ratio for Kv2.1 or Kv2.2 and Kv1.5, Kv4.2, or BK by Mann-Whitney (p<0.0001). C. Pearson correlation coefficients between GxTX-594 and GFP. Same cells as panel B. Significant differences were observed between correlation coefficients for Kv2.1 or Kv2.2 and Kv1.5, Kv4.2, or BK by Mann-Whitney (p<0.0001).

To quantify the relative labeling of the GFP-tagged K+ channels we measured the ratio of GxTX-594 to GFP fluorescence. CHO cells expressing the GFP-tagged variants of Kv2.1, Kv2.2, Kv4.2 + KChIP2, Kv1.5 + Kvβ2, and BK were incubated with 100 nM GxTX-594 for 5 minutes followed by washout with external solution to minimize background fluorescence (Fig 10 A). After background subtraction, the ratio of fluorescence excited at 594 nm (GxTX-594) to fluorescence excited at 488 nm (GFP) was normalized to the mean 594 nm to 488 nm ratio for Kv2.1-GFP. The 594:488 ratio was not distinguishable from zero for Kv4.2 + KChIP2, Kv1.5 + Kvβ2, or BK (Fig 10 B), indicating minimal, if any, binding to these channel subtypes. Furthermore, no colocalization was apparent between Kv4.2 + KChIP2, Kv1.5 + Kvβ2, or BK and the residual fluorescence at GxTX-594 wavelengths (Fig 10 C). A similar set of experiments conducted without the auxiliary subunits of Kv4.2 or Kv1.5 also gave no indication of GxTX-594 labeling of Kv4.2 or Kv1.5 (Fig Supplement 1 B, C, D). We did not conduct a comprehensive survey of the more than 80 known mammalian proteins containing voltage sensor domains, and thus cannot be certain that GxTX-594 does not bind any other voltage sensors, or other receptors on cell surfaces. However, the lack of labeling of these related voltage-gated K^+^ channels indicates that GxTX-594 does not promiscuously label voltage sensors. In all tests conducted, GxTX-594 was selective for Kv2 proteins.

To determine whether expression of Kv2 proteins embedded in tissue can be imaged with GxTX-594, we overexpressed Kv2.1-GFP in brain slices. We examined rat brain at CA1 pyramidal neurons of the hippocampus. We chose CA1 neurons for several reasons: they express Kv2 channels at a density typical of central neurons, the physiology of these neurons have been intensively studied, and their electrical properties are relatively homogeneous (Misonou, Mohapatra et al. 2005). Organotypic hippocampal slice cultures prepared from postnatal day 5-7 rats were sparsely transfected with Kv2.1-GFP, resulting in a subset of neurons displaying green fluorescence. When imaged two to four days after transfection, GFP fluorescence was observed in the plasma membrane surrounding neuronal cell bodies and proximal dendrites (Fig Supplement 4 A and B). Six days or more after transfection, Kv2.1-GFP fluorescence organized into clusters on the surface of the cell soma and proximal processes (Fig 11 A, Fig Supplement 4 C), a pattern consistent with a prior report of endogenous Kv2.1 in CA1 neurons (Misonou, Mohapatra et al. 2005). After identifying a neuron expressing Kv2.1-GFP, solution flow into the imaging chamber was stopped and GxTX-594 was added to the static bath solution to a final concentration of 100 nM. After five minutes of incubation, solution flow was restarted, leading to wash-out of excess GxTX-594 from the imaging chamber. After wash-out, GxTX-594 fluorescence remained colocalized with Kv2.1-GFP (Fig 11, Fig Supplement 4), indicating that GxTX-594 is able to permeate through dense neural tissue and bind to Kv2 proteins on neuronal surfaces. Pearson correlation coefficients confirmed the colocalization of GxTX-594 with Kv2.1-GFP in multiple slices (Fig 11 C). In most images of Kv2.1-GFP expressing neurons, GxTX-594 also labeled puncta on neighboring neurons that did not express Kv2.1-GFP, but at intensities that were roughly an order of magnitude dimmer (Fig 11 B, white arrow). The clustered GxTX-594 fluorescence patterns on the cell body, and proximal processes of CA1 neurons was strikingly similar to reported patterns of anti-Kv2 immunofluorescence patterns and is consistent with GxTX-594 labeling endogenous Kv2 proteins in CA1 neurons. While we cannot exclude the possibility that CA1 neurons have a subset of Kv2 proteins on their surface that are not labeled by GxTX-594, we saw no indication of Kv2.1-GFP on neuronal surfaces that are not labeled by GxTX-594.

**Figure 11:**
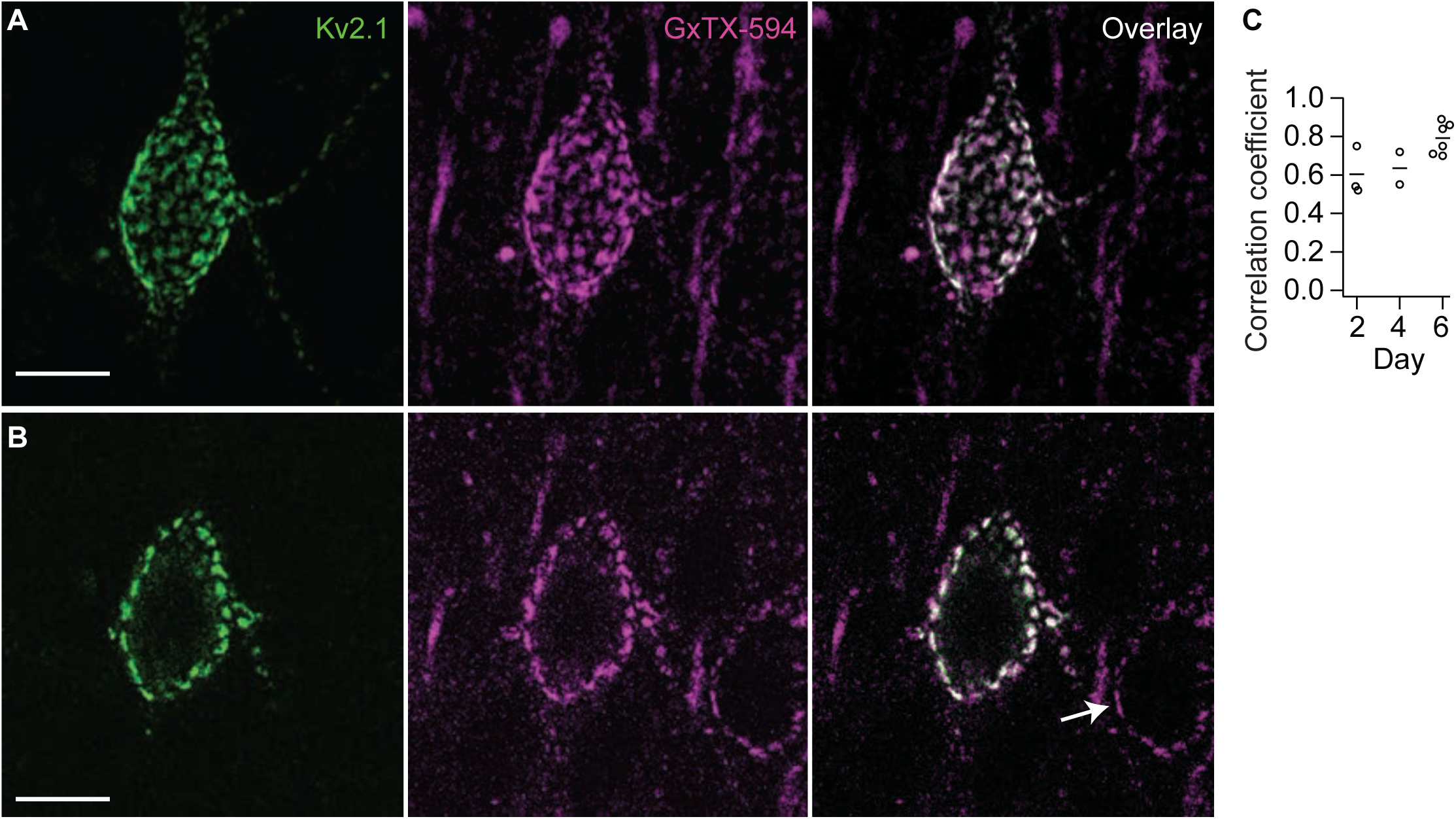
GxTX-594 labels CA1 hippocampal pyramidal neurons transfected with Kv2.1-GFP. A. Two-photon excitation images of fluorescence from the soma and proximal dendrites of a rat CA1 hippocampal pyramidal neuron in a brain slice six days after transfection with Kv2.1-GFP (Left), labeled with 100 nM GxTX-594 (Middle) and overlay (Right). Image represents a Z-projection of 20 optical sections. Scale bar 10 µm. B. A single optical section of the two-photon excitation image shown in A. GxTX-594 labels both Kv2.1-GFP puncta from a transfected cell and apparent endogenous Kv2 proteins from an untransfected cell in the same cultured slice (Right, arrow). C. Pearson correlation coefficients from CA1 hippocampal neurons two, four, and six days after transfection with Kv2.1-GFP. Each circle represents a different neuron. Bars are arithmetic means.

While we observed GxTX-594 fluorescence that morphologically resembles endogenous Kv2 protein localizations, we also found that GxTX-594 occasionally labels structures not consistent with Kv2 proteins (Fig Supplement 4 lower panel arrows). This non-Kv2 labeling was most prevalent at the surface of the hippocampal slices and progressively decreased as the imaging plane was moved deeper into the tissue (data not shown). Our interpretation of this phenomenon is that GxTX-594 can accumulate in the dead tissue and debris that is present at the surface of a hippocampal section after it is cut. For this reason, we analyzed only GxTX-594 fluorescence with subcellular localizations consistent with Kv2 channels.

### GxTX-594 labeling of neurons in brain slice responds to neuronal depolarization

To test whether GxTX-594 labeling of neurons in brain slices is consistent with labeling of endogenous Kv2 voltage sensors, we determined whether GxTX-594 labeling responds to voltage changes in tissue. First, we looked for Kv2-like labeling patterns on CA1 pyramidal neurons in untransfected brain slices bathed in 100 nM GxTX-594. With two-photon excitation, optical sections are thinner than the neuronal cell bodies, and fluorescence appeared as puncta circumscribed by dark intracellular spaces (Fig 12 A). This was similar to the patterns of fluorescence in Kv2.1-GFP transfected slices (Fig 11 B), and consistent with the punctate expression pattern of Kv2.1 in CA1 pyramidal neurons seen in fixed brain slices (Misonou, Mohapatra et al. 2005).

**Figure 12:**
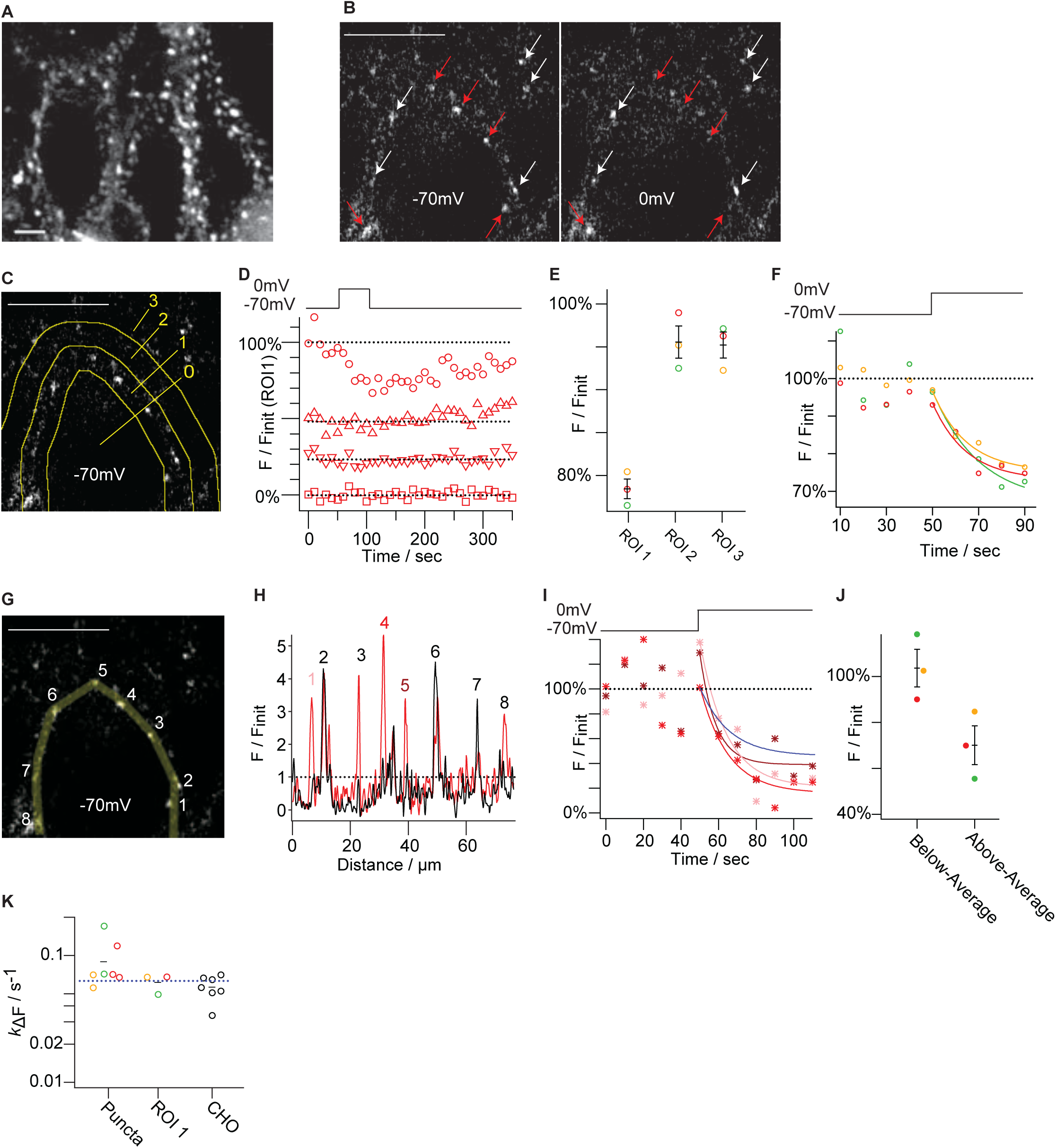
GxTX-594 puncta on hippocampal CA1 neurons are sensitive to voltage stimulus. A. Two-photon excitation optical section from CA1 pyramidal neurons in a cultured hippocampal slice after incubation with 100 nM GxTX-594. Scale bar is 5 µm. B. 594 fluorescence from a single two-photon excitation optical section before and during depolarization of a whole cell patch-clamped neuron. Text labels of holding potential (−70 mV or 0 mV) indicate approximate position of the patch-clamp pipette. Red arrows indicate stimulus-sensitive puncta, white arrows indicate stimulus-insensitive puncta. Left panel is the average fluorescence of the three frames at −70 mV before depolarization while the right panel is the average fluorescence of three frames after holding potential was stepped to 0 mV. Scale bar is 5 µm. Supplementary Movie 1 contains time-lapse images from this experiment. C. ROIs used in analysis for panels D, E, and F. Same slice as panel B. ROI 1 contains the apparent plasma membrane of the cell body of the patch-clamped neuron, it was generated by drawing a path tracing the apparent plasma membrane and then expanding to an ROI containing 15 pixels on either side of the path (1.2 µm total width). ROI 2 contains the area 15-45 pixels outside the membrane path (1.2 µm total width). RO1 3 contains the area more than 45 pixels outside of the membrane path. ROI 0 contains the area more than 15 pixels inside the membrane path. Scale bar is 5 µm. D. Fluorescence from each ROI shown in C. Squares represent ROI 0, circles represent ROI 1, upward triangles represent ROI 2 and upside-down triangles represent ROI 3. Background was defined as the mean fluorescence of ROI 0 during the experiment. *F_init_* was defined as the mean fluorescence of ROI 1 during the first six frames, after subtraction of background. Dotted lines represent the average fluorescence of the first six frames of each ROI. The voltage protocol is shown above. E. Change in fluorescence during a 0 mV step for different ROIs in multiple hippocampal slices. ROIs for each slice were defined by methods described in panel C. Circles represent mean fluorescence from six frames during 0 mV stimulus (*F*), normalized to mean fluorescence from same ROI in six frames before stimulus (*F_init_*). Circle color is consistent between ROIs for each hippocampal slice, red circles are from the slice shown in panel D. Black bars are arithmetic mean ± standard error from 3 hippocampal slices. F. Kinetics of fluorescence change from ROI 1 during a 0 mV step in multiple hippocampal slices. ROI 1 for each slice was defined by the method described in panel C. Lines are monoexponential fits (Eq.1). *k*_Δ*F*_ = 6.8 x 10^-2^ ± 2.6 x 10^-2^ s^-1^ (yellow), 6.8 x 10^-2^ ± 4.6 x 10^-2^ s^-1^ (red), and 4.9 x 10^-2^ ± 2.1 x 10^-2^ s^-1^ (green). The voltage protocol is shown above. Colors of individual circles indicate the same slices as panel E. G. Region of interest used in analysis for panels H, I, and J. Same image as panel C. The shaded ROI was generated by drawing a path tracing the apparent plasma membrane and then expanding to an ROI containing 5 pixels on either side of the path (0.4 µm total width). Numbers indicate puncta that appear as peaks in panel H. Scale bar is 5 µm. H. A plot of the fluorescence intensity along the ROI shown in panel G before (red trace) and during (black trace) 0 mV stimulus. Numbers above peaks correspond to puncta labeled in panel G. Red trace: mean fluorescence intensity during the three frames immediately before the stimulus, normalized to mean intensity of entire ROI (black dotted line), plotted against distance along path. Black trace: mean fluorescence intensity during three frames at 0 mV, normalized by the same *F_init_* value as the red trace. I. Kinetics of fluorescence change of individual puncta from panel G. Puncta intensity are average intensity of points extending to half maxima of each peak in panel H. Asterisks indicate mean fluorescence intensity of puncta 1 (pink), 4 (red), and 5 (dark red). Lines are monoexponential fits (Eq.1). *k*_Δ*F*_ = 7.1 x 10^-2^ ± 2.9 x 10^-2^ s^-1^ (pink), 6.7 x 10^-2^ ± 2.5 x 10^-2^ s^-1^ (red) and 1.2 x 10^-1^ ± 5.6 x 10^-2^ s^-1^ (dark red). Fits to other puncta had standard deviations larger than *k*_Δ*F*_ values, and were excluded. *F_init_* was defined as the mean background subtracted fluorescence of the puncta during the six frames before stimuli. The voltage protocol is displayed above. Blue line is prediction from Scheme I. J. Comparison of fluorescence change of puncta and inter-puncta regions in response to 0 mV stimulus. The regions before stimulus shown in the red line of panel H that had *F/F_init_* ≥ 1 (above average) were binned separately from regions with *F/F_init_* < 1. The mean fluorescence of each region during 0 mV stimulus (panel H black trace) was compared to the fluorescence before stimulus (panel H red trace). Circles indicate values from three independent hippocampal slices, colors indicate same slices as panel E and F. Black bars are arithmetic mean ± standard error. K. Rate of fluorescence change of GxTX-594 after a 0 mV stimulus; from puncta (as in panel I), ROI 1 (as in panel F) or Kv2.1-CHO cells in 100 nM GxTX-594. Kv2.1-CHO cells were imaged at the same temperature as neurons (30 °C) using neuronal intracellular solution. CHO extracellular solution (CE) was used for Kv2.1-CHO cell experiments. Kv2.1-CHO measurements were made by airy disk confocal imaging. Black bars are mean ± SEM from data shown. Dashed blue line is *k*_Δ*F*_ = 6.31 x 10^-2^ s^-1^ prediction of Scheme I.

We tested whether the punctate fluorescence was voltage-sensitive using voltage clamp. To ensure voltage clamp of the neuronal cell body, slices were bathed in tetrodotoxin to block Na+ channels, and Cs+ was included in the patch pipette solution to block K+ currents. In each experiment, whole-cell configuration was achieved with a GxTX-594 labeled neuron, and holding potential was set to −70 mV. At this point, time-lapse imaging of a two-photon excitation optical section was initiated. Depolarization to 0 mV resulted in loss of fluorescence from a subset of puncta at the perimeter of the voltage-clamped neuron (Fig 12 B red arrows, Supplemental Movie). Other fluorescent puncta appeared unaltered by the 0 mV step (Fig 12 B, white arrows). These puncta could represent Kv2 proteins on a neighboring cell, off-target labeling by GxTX-594, or voltage-insensitive Kv2 proteins, possibly due to near-surface internalization. To assess if the fluorescence changes were due to the voltage change or random fluctuations during imaging, we compared fluorescence of the apparent cell membrane region to regions more distal from the voltage-clamped cell body. To quantify this fluorescence change, an ROI 30 pixels (1.2 µm) wide containing the apparent membrane of the cell body was compared to other regions within each image (Fig 12 C). The region containing the membrane of the cell body lost fluorescence during the 0 mV step (Fig 12 D, ROI 1) while neither of the regions more distal (ROI 2, ROI 3), nor the intracellular region (ROI 0) showed a similar change. In multiple slices, the voltage-dependent decrease of fluorescence was seen specifically in the region of the voltage-clamped neuronal membrane (Fig 12 E). The kinetics of fluorescence response of the voltage-clamped membrane region were similar between slices (Fig 12 F, K).

To determine whether the fluorescence response to depolarization was driven by the Kv2-like puncta on the cell membrane, the fluorescence along a path containing the apparent cell membrane was selected by drawing a path connecting fluorescent puncta surrounding the dark cell body region, and averaging fluorescence in a region 10 pixels wide (0.4 µm) centered on this path (Fig 12 G, yellow line). The fluorescence intensity along the path of the ROI revealed distinct peaks corresponding to puncta (Fig 12 H, red trace). After stepping the neuron to 0 mV, the intensity of fluorescence of a subset of puncta lessened (Fig 12 H, black trace, peaks 1,3,4,5,8). While the data is quite noisy, it is clear that even more reduction is observed in individual fluorescence puncta in brain slice (Fig 12, I blue line). This suggests the voltages sensors in these functionally-selected puncta are extensively activated. The EVAP model predicts only a 55% reduction of fluorescence from CHO cells stepped to 0 mV in 100 nM GxTX. However, the GxTX-594 concentration within tissue may be more dilute than in the bath solution, and consequently the sensitive fraction calculated from the EVAP model is a lower bound. Other puncta maintained or increased their brightness after depolarization (Fig 12 H, black trace, peaks 2, 6, 7), but it was not certain whether these voltage-insensitive puncta correspond to Kv2 proteins on the surface of the same voltage-clamped neuron. When the kinetics of fluorescence intensity decay of individual voltage-sensitive puncta were fit with Eq. 1, *k*_Δ*F*_ values were similar between puncta (Fig 12 I), consistent with these spatially separated puncta all being on the surface of the voltage-clamped neuron. The fluorescence changes measured in these puncta were similar to those predicted by the Fig 8 model (Fig 12 I, blue line), although the variability of these measurements was substantial. To address whether the fluorescence change at the cell membrane was driven by decreases in regions of punctate fluorescence, the punctate and non-punctate fluorescence intensity changes were analyzed separately. The regions with fluorescence intensities above average for the path (Fig 12 H, dashed line) were binned as one group. The above-average group, by definition, contained all punctate fluorescence. When comparing the fluorescence before and during the 0 mV step, the regions that were initially below average fluorescence maintained the same intensity (103 ± 8%); regions that were initially above average fluorescence decreased in intensity (70 ± 8%) (Fig 12 J). This suggests that the detectable unlabeling along the cell membrane originated in the Kv2-like puncta, not the regions of lower fluorescence intensity between them.

To determine whether the dynamics of the voltage-dependent responses of GxTX-594 fluorescence on neurons in brain slices were consistent with labeling of Kv2.1, we performed experiments with Kv2.1-expressing CHO cells under similar conditions as brain slice: 100 nM GxTX-594, 30°C, Cs^+^-containing patch pipette solution. The rate of fluorescence changes in Kv2.1-CHO cells were similar to neurons in brain slices, and consistent with the *k*_Δ*F*_ of 0.06 s^-1^ predicted by Fig 8 model (Fig 12 K). However, the data underlying of *k*_Δ*F*_ measurements was noisy, limiting our ability to detect differences.

## Discussion

The molecular targeting, conformation selectivity, and spatial precision of fluorescence from GxTX-594 enable identification of where in tissue the conformational status of Kv2 voltage sensors becomes altered. However, the utility of GxTX-594 as an EVAP is limited by several factors including emission intensity, variability between experiments, and inhibition of Kv2 proteins. We discuss the potential utility and limitations of the EVAP mechanism underlying GxTX-594.

### Unique capabilities of GxTX-594

The Kv2 EVAP presented here is the only imaging method we are aware of for measuring voltage-sensitive conformational changes of a specific, endogenous protein. As GxTX binding selectively stabilizes the fully resting conformation of Kv2.1 voltage sensors (Tilley, Angueyra et al. 2019), GxTX-594 labeling is expected to report the fully resting conformation of the Kv2 voltage sensor, in which the first gating charge of the Kv2 S4 segment is in the gating charge transfer center (Tao, Lee et al. 2010). Images of GxTX-594 fluorescence reveal this conformation’s occurrence, with subcellular spatial resolution. Importantly, the EVAP model we develop allows deconvolution of the behavior of unlabeled Kv2 proteins from Kv2 proteins inhibited by GxTX-594. This enables the subcellular locations where Kv2 voltage sensing occurs to be seen for the first time.

Electrophysiological approaches can detect the voltage-sensitive K^+^ conductance of Kv2 channels. However, the majority of Kv2 proteins on cell surface membranes do not function as channels and are nonconducting (Benndorf, Koopmann et al. 1994, O’Connell, Loftus et al. 2010). Kv2 proteins dynamically regulate cellular physiology by nonconducting functions (Antonucci, Lim et al. 2001, Singer-Lahat, Sheinin et al. 2007, Feinshreiber, Singer-Lahat et al. 2010, Dai, Manning Fox et al. 2012, Fox, Haberkorn et al. 2015, Johnson, Leek et al. 2018, Kirmiz, Palacio et al. 2018, Kirmiz, Vierra et al. 2018, Kirmiz, Gillies et al. 2019, Vierra, Kirmiz et al. 2019). The Kv2 EVAP reports on the conformation of Kv2 voltage sensors independently from ion conductance, enabling the study of voltage sensor involvement in Kv2’s nonconducting physiological functions. We present this EVAP as a prototype molecular probe for imaging voltage sensing by endogenous proteins, in tissue, with molecular specificity.

Here, during our initial testing of GxTX-594, we observed that the majority of Kv2 protein detected at discrete individual clusters was voltage sensitive. While it may not sound surprising to find that voltage-gated ion channel proteins are voltage sensitive, the voltage sensors of surface-expressed proteins can be immobilized. For example, gating charge of the L-type Ca^2+^ channel Cav1.2 is immobilized until it is bound by an intracellular protein (Turner, Anderson et al. 2020). Kv2.1 channel function is extensively regulated by neurons. In rat CA1 neurons the clustered Kv2 channels are proposed to be nonconducting (Fox, Loftus et al. 2013). Our results show that clustered Kv2 proteins in rat CA1 neurons remain voltage-sensitive. Interestingly, when Kv2.1 is expressed in CHO cells, a fraction of the GxTX-594 fluorescence is voltage insensitive (Fig 7 C). This observation is consistent with voltage sensor immobilization of some surface-expressed Kv2.1 protein, although it could be due to intracellular Kv2.1–GxTX-594 proteins that appear to be at the cell surface.

We used images of GxTX-594 fluorescence to measure the coupling between endogenous Kv2 proteins and membrane potential at specific, subcellular anatomical locations. Similarly, GxTX-594 imaging should detect changes of voltage sensor status when Kv2 proteins become engaged or disengaged from nonconducting functions such as formation of plasma membrane– endoplasmic reticulum junctions (Antonucci, Lim et al. 2001, Fox, Haberkorn et al. 2015, Kirmiz, Palacio et al. 2018), regulation of exocytosis (Singer-Lahat, Sheinin et al. 2007, Feinshreiber, Singer-Lahat et al. 2010), regulation of insulin secretion (Dai, Manning Fox et al. 2012) interaction with kinases, phosphatases and SUMOylases (Misonou, Mohapatra et al. 2004, Park, Mohapatra et al. 2006, Dai, Kolic et al. 2009, Cerda and Trimmer 2011, McCord and Aizenman 2013), formation of specialized subcellular calcium signaling domains (Vierra, Kirmiz et al. 2019), and interactions with astrocytic end feet (Du, Tao-Cheng et al. 1998). GxTX-594 could potentially reveal conformational changes in organs throughout the body where Kv2 proteins are expressed, which include muscle, thymus, spleen, kidney, adrenal gland, pancreas, lung, and reproductive organs (Bocksteins 2016).

### Limitations of GxTX-594

There are important limitations to the GxTX-594 approach, and of the underlying EVAP mechanism generally. We discuss several limitations which are worth considering in the design of any studies.

#### GxTX-594 is slow

The kinetics of GxTX-594 labeling are limited to measuring changes in Kv2 activity on a time scale of tens of seconds. Thus GxTX-594 will not provide sufficient time resolution to distinguish fast electrical signaling events. However, we have found that the temporal resolution of GxTX-594 is compatible with live imaging and electrophysiology experiments. The response time of GxTX-594 is far slower than the kinetics of Kv2 conformational change, limiting measurements to the probability, averaged over time, that voltage sensors are resting or active. It is worth noting that the probability of a conformation’s occurrence is a valuable measure, and the ultimate quantitation of many biophysical studies of ion channels (e.g., open probability, steady state conductance, gating charge-voltage relation).

#### GxTX-594 dynamics are altered in confined extracellular spaces

The kinetics of GxTX-594 dynamics varied within different regions of the same CHO cell. GxTX-594 fluorescence in the restricted space at the center of the glass-adhered surface appeared to respond more slowly to voltage changes than the periphery, which has better access to the bath solution (Fig 6). The location dependence of *k*_Δ*F*_ was more pronounced during GxTX-594 labeling at −80 mV than unlabeling at +40 mV, and such a difference can be explained by the distinct voltage dependent affinities of Kv2.1 for GxTX-594. We suspect that the more extreme location dependence at −80 mV is due to a high density of Kv2.1 binding sites in the restricted extracellular space between the cell membrane and glass coverslip, such that GxTX-594 is depleted from solution by binding Kv2.1 before reaching the center of the cell. After unbinding at +40 mV, each GxTX-594 molecule is expected to be less likely to rebind to Kv2.1 proteins which are in an activated, low affinity conformation (Tilley, Eum et al. 2014, Tilley, Angueyra et al. 2019).

The different GxTX-594 dynamics in a restricted space is significant because the space extracellular to Kv2.1 channel clusters is restricted in many neurons. The Kv2 channel clusters in the plasma membrane of hippocampal and cortical interneurons are tightly associated with astrocytic end feet and the cleft between these two membranes is a few nanometers wide (Du, Tao-Cheng et al. 1998). Experiments investigating Kv2 channels localized to these subcellular regions should consider the limitations of using EVAP probes in restricted spaces; most notably the finding that labeling dynamics are more dramatically affected than unlabeling dynamics (Fig 6 D and E).

#### GxTX-594 dynamics are variable

The GxTX-594 response rates and amplitudes were notably variable between Kv2.1 CHO cells. We did not succeed in constraining this variability and offer conjecture about its origins. Some variability results from fits of *k*_Δ*F*_ where the final value of the relaxation was poorly determined by the data (Fig 3C, 7B, 9B and Fig Supplement 2). Fluctuations in temperature during imaging likely had only a small effect on the fluorescence dynamics of GxTX-594; considering the measured Q_10_ of 3.8, the 2°*C* temperature fluctuations between the experiments in Figures 7 and 9 should produce only 1.3-fold change in *k*_Δ*F*_. We also expect that changes in fluorescence due to photobleaching of the Alexa Flour 594 fluorophore were limited, based on the observation that fluorescence remained unchanged in cells that were not voltage clamped but in the same field of view as voltage clamped cells receiving voltage stimulus (Fig 9 A).

We noticed that results were more consistent between stimuli of the same cell and the variability was greatest between cells. We suspect that cell-to-cell differences in Kv2.1 conformational equilibria are responsible for much of the variability in GxTX-594 response. The Kv2.1 conductance-voltage relation is regulated by many cellular pathways including kinases, phosphatases, and SUMOylases (Misonou, Mohapatra et al. 2004, Park, Mohapatra et al. 2006, Dai, Kolic et al. 2009, Cerda and Trimmer 2011, McCord and Aizenman 2013). Large cell-to-cell variation in Kv2.1 conductance-voltage, and gating charge-voltage relations have been reported in CHO cells (McCrossan, Roepke et al. 2009, Tilley, Eum et al. 2014, Kang, Vanoye et al. 2019, Tilley, Angueyra et al. 2019).

In this study, when we predicted the voltage sensor V_1/2_ of Kv2.1 from electrophysiology we observed a 6.4 mV standard deviation with a range of 19 mV, and this variance appeared to be exacerbated by GxTX-594 (Fig 5E) (9.7 mV standard deviation and range of 36 mV). As EVAP dynamics and the GV are both determined by voltage sensor activation, variability in the GxTX-594 response is expected. The hypothesis that cell-to-cell variation in fluorescence dynamics is due to the inherent variability of Kv2.1 voltage sensor activation could be more definitively tested by identifying if a correlation exists between the *V_1/2_* of the QV and fluorescence–voltage relationship from individual cells labeled with GxTX-594. While we have not attempted this, the structure of the variance in GxTX-594 fluorescence–voltage relationships is informative. The fluorescence–voltage relationships compiled from many cells become more variable near the midpoint of relevant voltage sensor movements. The response amplitude, *F/F_init_*, (Fig 7 D), is determined by unlabeled voltage sensor activation, and appears most variable near the *V_1/2_* of the unlabeled QV relation, −32 mV (Tilley, Angueyra et al. 2019). The response kinetics, *k*_Δ*F*_, are determined by activation of labeled voltage sensors and appear to become increasingly variable at voltages higher than −20 mV (Fig 7 E bottom panel). Despite the increasing variability, the *V_1/2_* and *z* from the Boltzmann fit of the *k*_Δ*F*_–voltage relationship to measurements from many cells was remarkably close to the same measurements in the QV of the GxTX–Kv2.1 complex with a *V_1/2_* and *z* that differ by 3.3 mV and 0.07 *e*_0_ respectively.

Another possibility is that variability in surface membrane composition undergirds the variability of the GxTX-594 response. GxTX affinities are influenced by the dynamically changing complement of lipids in the plasma membrane lipid composition. Lipids bind to the voltage sensing domain near the GxTX binding site (Milescu, Bosmans et al. 2009, Gupta, Zamanian et al. 2015). Sphingomyelinase D treatment, which alters the membrane composition by converting sphingomyelin to ceramide 1-phosphate, has been shown to enhance GxTX affinity for Kv2.1 by 4-fold (Milescu, Bosmans et al. 2009). Voltage activation of Kv2.1 is also affected by sphingomyelinase treatments. While variation in lipid composition is expected to cause variation in GxTX-594 dynamics, we do not know the degree to which the lipid composition varies between the Kv2.1-CHO cells in our studies.

#### Signal Intensity and Noise

Fluorescence signal intensity limits interpretation of EVAP labeling. We found that variability in the intensity of GxTX-594 labeling of CA1 neurons limited the quantitative interpretation with the EVAP model. However, the signal to noise was clearly sufficient to identify voltage sensing of endogenous Kv2 proteins. As CA1 hippocampal neurons express Kv2 proteins at a density typical of central neurons (Misonou, Mohapatra et al. 2004, Vacher, Mohapatra et al. 2008, Speca, Ogata et al. 2014). Thus, we expect that GxTX-594 imaging could be implemented in most brain regions with similar signal to noise characteristics. In other cell types, such as neurons of the subiculum, or the inner segment of photoreceptors, Kv2 proteins are expressed at far higher densities (Maletic-Savatic, Lenn et al. 1995, Gayet-Primo, Yaeger et al. 2018). Improved signal to noise would be expected from such cells that express higher densities of Kv2 proteins. Kv2 proteins are also expressed by many other cell types throughout the body (Bocksteins 2016). It remains to be seen what the signal to noise characteristics are for EVAP Kv2 labeling in these other tissues.

#### GxTX-594 inhibits Kv2 proteins

GxTX-based probes inhibit the Kv2 proteins they label by stabilizing the resting conformation of Kv2 voltage sensors. The Kv2.1–GxTX-594 complex does not open to conduct K^+^ ions in the physiological voltage range (Fig 5). Thus, GxTX-594 depletes the population of Kv2 proteins responding normally to physiological stimuli. While the concentration of an EVAP can be lowered to minimize interference with Kv2 activity, fluorescence intensity is diminished and further limits the ability to detect subtle physiological changes. With a related probe, we explored the impact of decreasing toxin concentration on fluorescence response of an EVAP (Tilley, Eum et al. 2014). Here, we demonstrate that lower concentration and physiological stimuli are not always required for scientifically meaningful implementation of the probe. Even when using GxTX-594 concentrations that grossly inhibit most Kv2 proteins, the behavior of unlabeled Kv2 proteins can be calculated using EVAP model we have developed. Of course, the electrical feedback within cells will be altered by such protocols.

#### The EVAP model is oversimplified

Another limitation of the analysis developed here is that the model of Kv2 voltage sensor conformational change is an oversimplification. The gating dynamics of Kv2 channels are more complex than our model (Islas and Sigworth 1999, Scholle, Dugarmaa et al. 2004, Jara-Oseguera, Ishida et al. 2011, Tilley, Angueyra et al. 2019). Under some conditions the assumption of voltage sensor independence will limit the model’s predictive power. Additionally, the model of GxTX-594 labeling developed here assumes that voltage sensors are in continuous equilibrium. These deviations from equilibrium likely explain the deviation of the model from the data in response to high frequency voltage steps (Fig 9 C, D).

### Conformation-selective probes reveal conformational changes of endogenous proteins

Measurements of dynamic labeling by a conformation-selective probe such as GxTX can enable deduction of how unlabeled proteins behave. This is perhaps counterintuitive because GxTX inhibits voltage sensor movement of the Kv2 protein it binds, and thus only bound proteins generate optical signals. This approach is analogous to calcium imaging experiments, which have been spectacularly informative about physiological calcium signaling (Yang and Yuste 2017), despite the fact that no optical signals originate from the physiologically relevant free Ca^2+^, only from Ca^2+^ that is chelated by a dye. In all such experiments, fluorescence from Ca^2+^-bound dyes is deconvolved using the statistical thermodynamics of Ca^2+^ binding to calculate free Ca^2+^ (Adams 2010). Similarly, GxTX-based probes dynamically bind to unlabeled Kv2 proteins, and the binding rate is dependent on the probability that unlabeled voltage sensors are in a resting conformation (Fig 8). Thus, the conformations of unlabeled Kv2 proteins influence the dynamics of labeling with GxTX-based probes. Consequently, the dynamics of labeling reveal the conformations of unlabeled Kv2 proteins.

Deployment of GxTX-594 to report conformational changes of endogenous proteins demonstrates that conformation-selective ligands can be used to image occurrence of the conformations they bind to. The same principles of action apply to any conformation-selective labeling reagent, suggesting that probes for conformational changes of many different proteins could be developed. Probes could conceivably be developed from the many other voltage sensor toxins and other gating modifiers that act by a similar mechanism as GxTX, yet target the voltage sensors of different ion channel proteins (McDonough, Mintz et al. 1997, Sack, Aldrich et al. 2004, Catterall, Cestele et al. 2007, Swartz 2007, Schmalhofer, Calhoun et al. 2008, Peretz, Pell et al. 2010, McCormack, Santos et al. 2013, Ahuja, Mukund et al. 2015, Dockendorff, Gandhi et al. 2018, Zhang, Sharma et al. 2018). Conformation-selective binders have been engineered for a variety of other proteins, and methods to quantify conformational changes from their fluorescence are needed. For example, fluorescently-labeled conformation-selective binders have revealed that endocytosed GPCRs continue to remain in a physiologically activated conformation (Irannejad, Tomshine et al. 2013, Tsvetanova, Irannejad et al. 2015, Eichel and von Zastrow 2018). A means to determine the conformational equilibria of GPCRs from fluorescence images has not yet been developed. We suggest that the statistical thermodynamic framework developed here could provide a starting point for more quantitative interpretation of other conformation-selective molecular probes.

## Acknowledgements

We thank Jim Trimmer (University of California, Davis) for numerous discussions, and constructive critical reading of an early version of the manuscript. We thank Georgeann Sack (Afferent LLC) for critical reading, editing, and feedback. This research was supported by US National Institutes of Health grants R01NS096317, U01NS090581, R21EY026449, and T32GM007377. P. Thapa was supported by American Heart Association postdoctoral fellowship 17POST33670698. GxTX variants were synthesized at the Molecular Foundry of the Lawrence Berkeley National Laboratory under US Department of Energy contract DE-AC02-05CH11231. The authors declare no competing financial interests.

## Author contributions (Credit nomenclature)

Parashar Thapa: Conceptualization, Formal analysis, Investigation, Methodology, Visualization, Writing-original draft, Writing- reviewing & editing

Robert Stewart: Conceptualization, Formal analysis, Investigation, Methodology, Visualization, Writing- original draft, Writing- reviewing & editing

Rebecka J. Sepela: Conceptualization, Formal analysis, Investigation, Methodology, Visualization, Writing- original draft, Writing- reviewing & editing

Oscar Vivas: Investigation, Methodology, Writing- reviewing & editing Laxmi K. Parajuli: Investigation, Methodology, Writing- reviewing & editing

Mark Lillya: Conceptualization, Formal analysis, Investigation, Methodology, Visualization Sebastian Fletcher-Taylor: Investigation, Methodology, Writing- reviewing & editing

Bruce E. Cohen: Conceptualization, Funding acquisition, Project administration, Supervision, Writing- reviewing & editing

Karen Zito: Conceptualization, Funding acquisition, Investigation, Methodology, Project administration, Supervision, Writing-original draft, Writing- reviewing & editing

Jon T. Sack: Conceptualization, Formal analysis, Funding acquisition, Investigation, Methodology, Project administration, Supervision, Visualization, Writing- original draft, Writing- reviewing & editing

---

**Figure Supplement 1:**
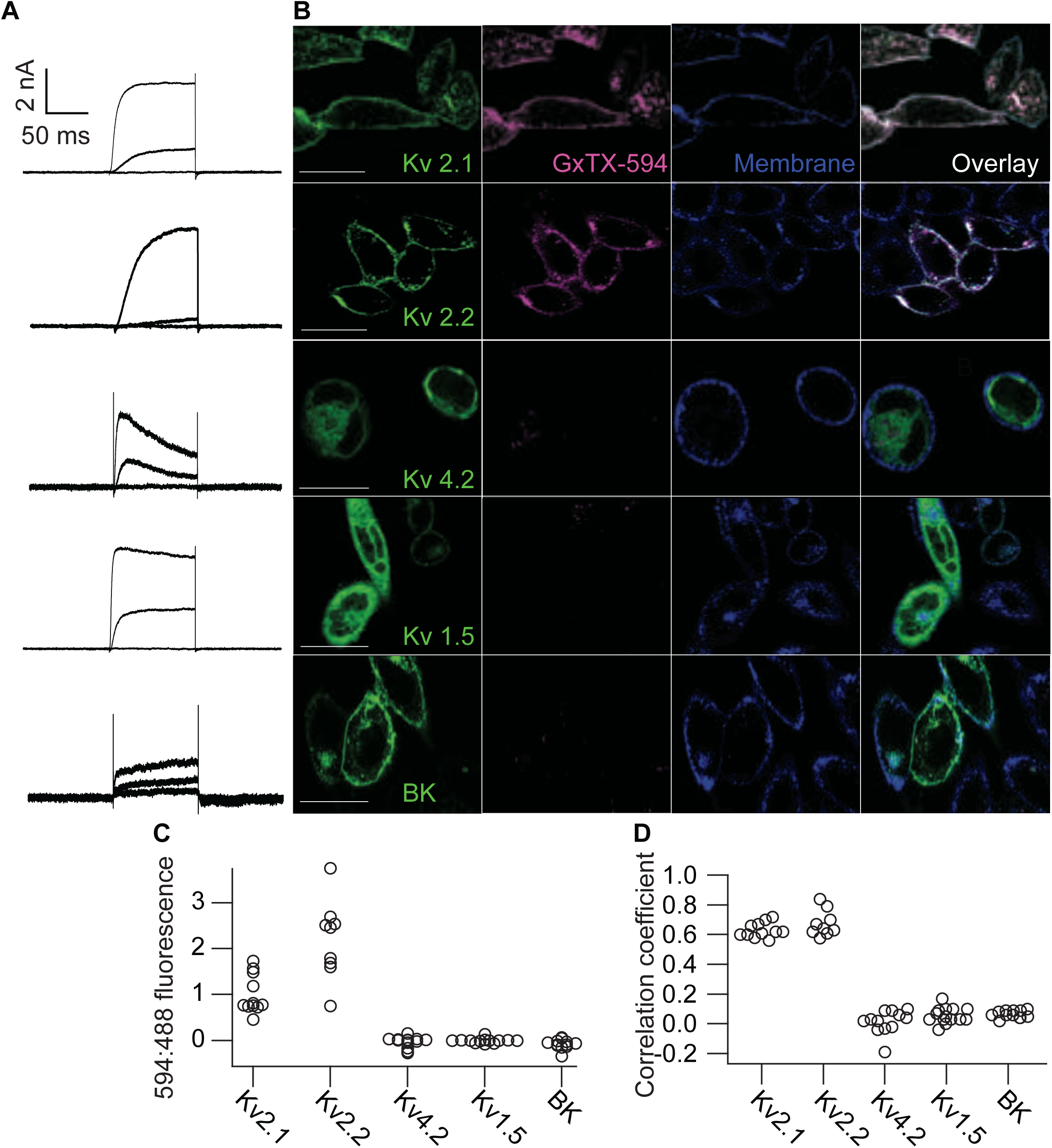
GxTX-594 selectively labels Kv2 proteins on cell surfaces. A. Exemplar whole-cell voltage clamp recordings of CHO cells expressing Kv2.1-GFP, Kv2.2-GFP, Kv4.2-GFP, Kv1.5-GFP, or BK-GFP. Recordings shown are representative responses to 100 ms steps from −100 mV to −40, 0, and +40 mV. B. Confocal imaging of fluorescence from live CHO cells transfected with Kv2.1-GFP, Kv2.2-GFP, Kv4.2-GFP Kv1.5-GFP, or BK-GFP (Indicated by row) and labeled with GxTX-594. Confocal imaging plane was above the glass-adhered surface. Cells were incubated with 100 nM GxTX-594 and 5 μg/mL WGA-405 and rinsed before imaging. Fluorescence shown corresponds to emission of GFP (Column 1), Alexa Fluor 594 (Column 2), WGA-405 (Column 3), or an overlay of GFP, Alexa Fluor 594, and WGA-405 (Column 4). Scale bars are 20 µm. C. Fluorescence intensity resulting from 594 nm excitation of GxTX-594 is divided by fluorescence intensity resulting from 488 nm excitation of GFP. Analysis methods as in Figure 10 B. Kv2.1 n = 11, Kv2.2 n = 9, Kv4.2 n = 13, Kv1.5 = 13, and BK n =12. Cells were compiled from two independent GxTX-594 applications, each circle corresponds to a cell. Significant differences were observed between GxTX:GFP ratio for Kv2.1 or Kv2.2 and Kv1.5, Kv4.2, or BK by Mann-Whitney (p<0.0001). D. Pearson correlation coefficients between GxTX-594 and GFP. Same cells as panel C. Analysis methods as in Figure 10 C.

**Figure Supplement 2:**
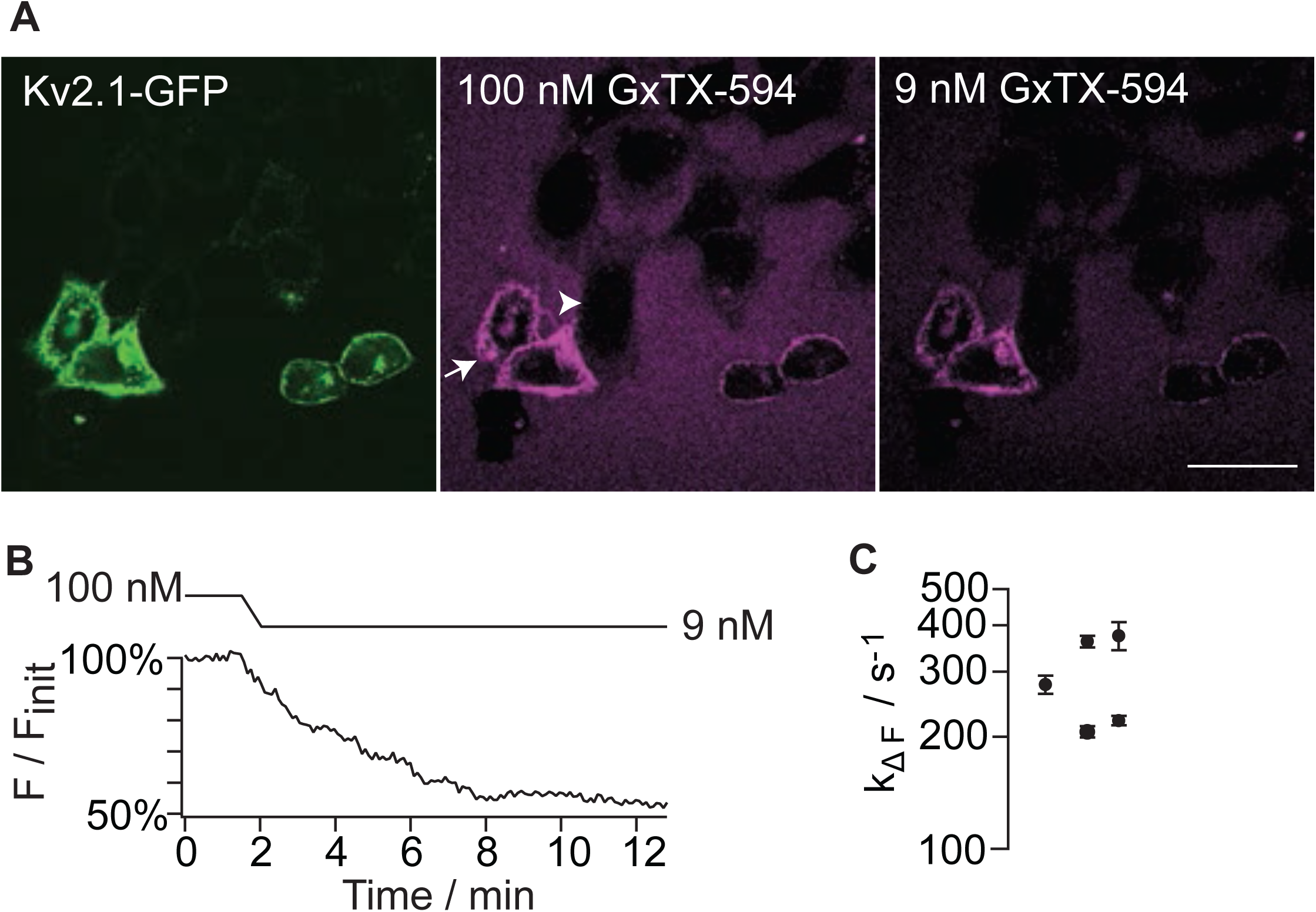
GxTX-594 fluorescence equilibration after dilution from 100 nM to 9nM. A. CHO cells transfected Kv2.1-GFP (Left) with in 100 nM GxTX-594 (Middle) and after dilution to 9 nM GxTX-594 (Right). Regions darker than the background bath solution are CHO cells not transfected with Kv2.1-GFP (Arrowhead). Scale bar is 20 μm. B. Fluorescence excited at 594 nM during dilution from 100 nM GxTX-594 to 9 nM GxTX-594. Timing of dilution is shown above the graph. Fluorescence was normalized to the mean 594 fluorescence before dilution to 9 nM. Data are from cell labeled with arrow in panel A. 100 nM and 9 nM images in panel A are from 0 and 12 minutes respectively. C. Rate of GxTX-594 fluorescence decay after dilution from monoexponential fits (Eq 1). Error bars are standard deviation of fit parameter. n = 5 cells.

**Figure Supplement 3:**
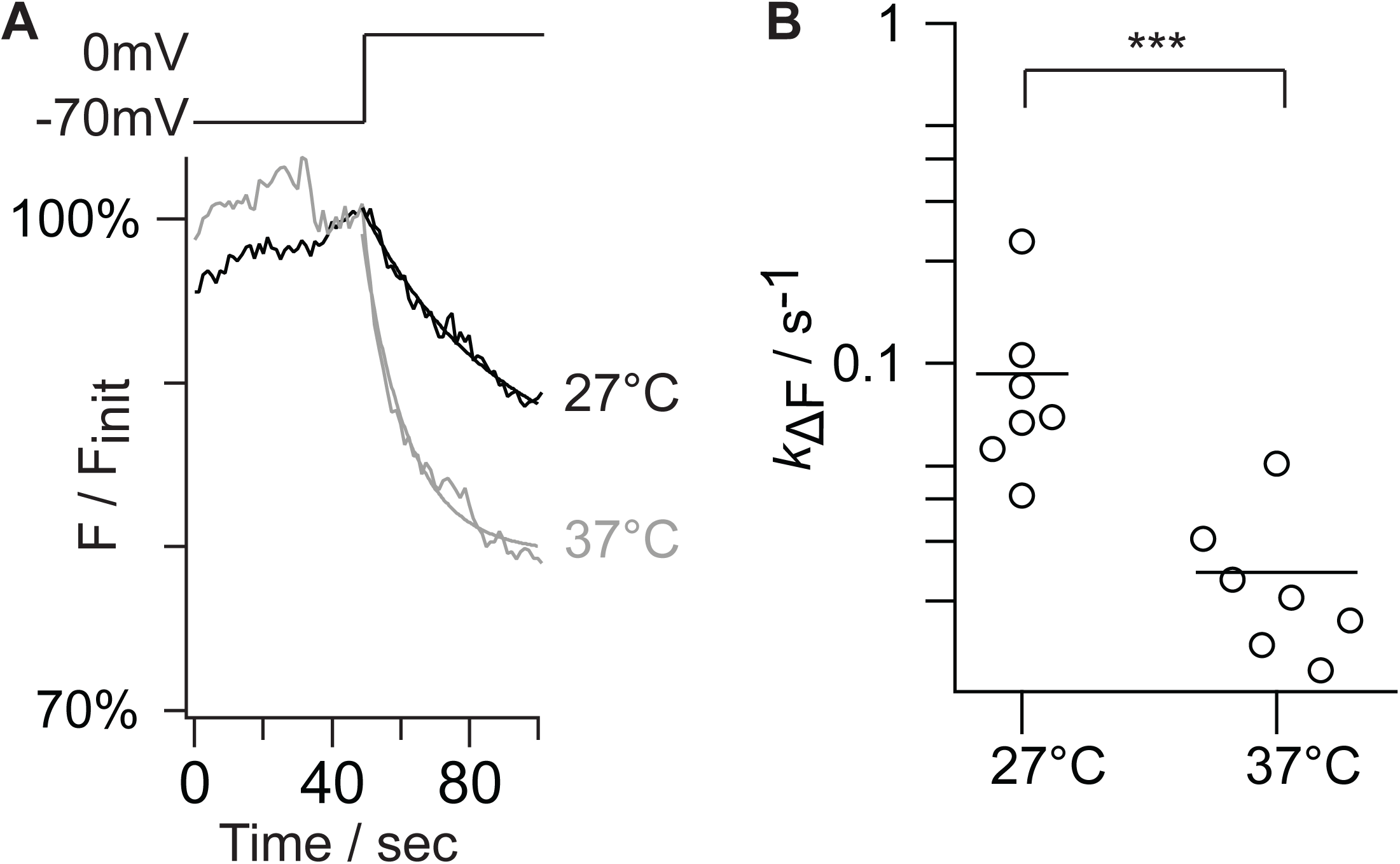
Variation in bath temperature does not account for variability of GxTX-594 kinetics. A. Representative traces of GxTX-594 fluorescence intensity response to voltage changes at 27 °C (black) and 37 °C (gray). Smooth lines are fits of a monoexponential function (Eq. 1): 27 °C *k*_Δ*F*_ = 2.47 x 10^-2^ ± 0.39 x 10^-2^ s^-1^; 37 °C *k*_Δ*F*_ = 7.54 x 10^-2^ ± 0.54 x 10^-2^ s^-1^. Background was determined by taking the average fluorescence of a region that did not contain cells over the time course of the voltage protocol. This average was subtracted from each experimental group. Traces were normalized to initial fluorescence intensity before the application of the voltage stimulus. B. *k*_Δ*F*_ at 27 °C and 37 °C. The rate of fluorescence change was significantly faster at higher temperatures (Mann-Whitney p = 0.0005). From geometric means (Bars), a Q_10_ of 3.8 was calculated between 27 °C and 37 °C. Each circle represents one cell, n = 7 both groups.

**Figure Supplement 4:**
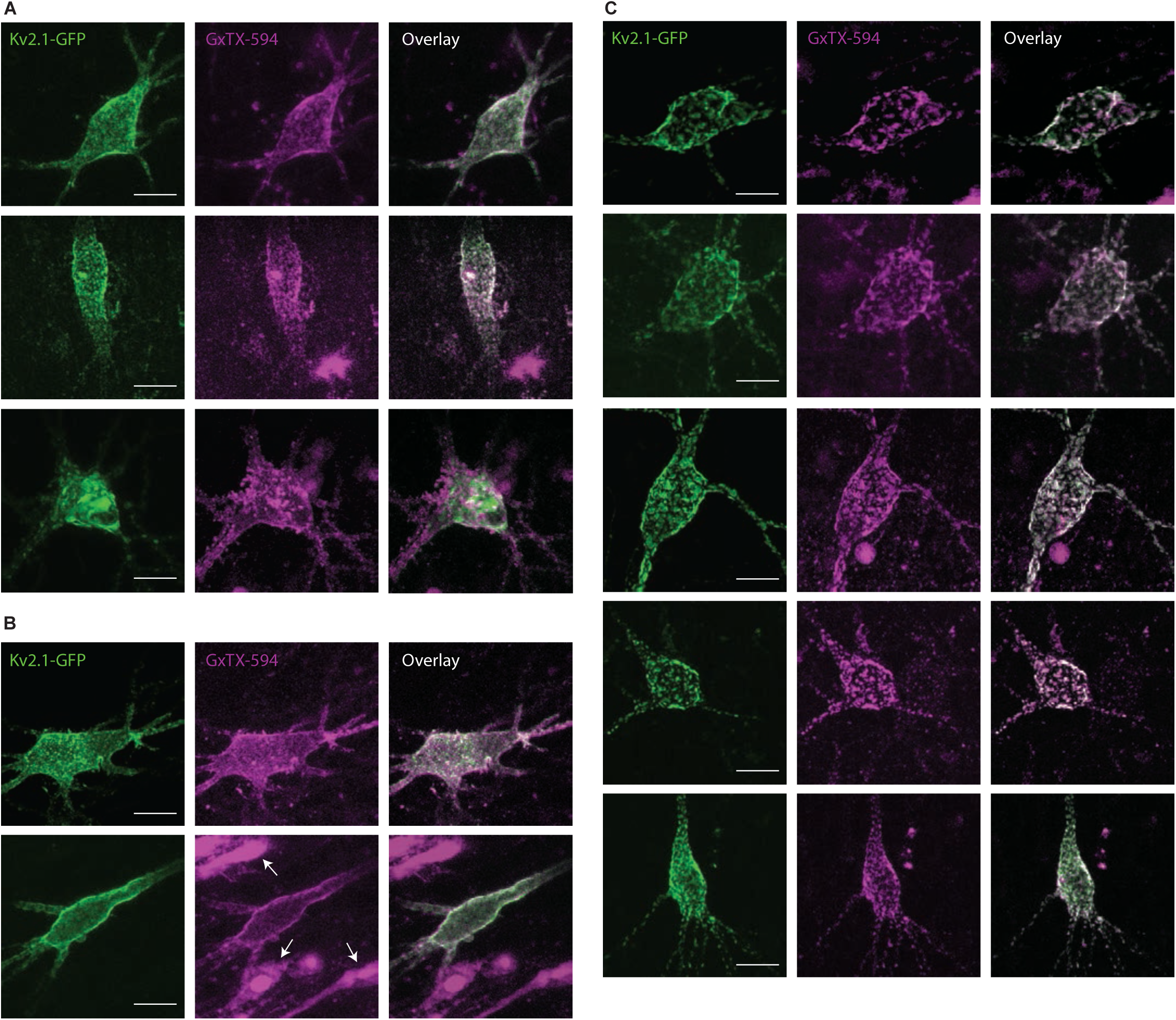
GxTX-594 labels CA1 hippocampal pyramidal neurons transfected with Kv2.1-GFP. Two-photon excitation images of rat CA1 hippocampal pyramidal neurons in brain slices as in Figure 7 A. Kv2.1-GFP (Left), GxTX-594 (Middle) and overlay (Right). Scale bars 10 µm. A. Pyramidal neurons two days after transfection with Kv2.1-GFP. B. Pyramidal neurons four days after transfection with Kv2.1-GFP. C. Pyramidal neurons six days after transfection with Kv2.1-GFP.

## Notes

### Competing Interest Statement

The authors have declared no competing interest.

## References

Adams, S. R. (2010). “How calcium indicators work.” Cold Spring Harb Protoc 2010(3): pdb.top70.

Aggarwal, S. K. and R. MacKinnon (1996). “Contribution of the S4 segment to gating charge in the Shaker K+ channel.” Neuron 16(6): 1169–1177.

Ahuja, S., S. Mukund, L. Deng, K. Khakh, E. Chang, H. Ho, S. Shriver, C. Young, S. Lin, J. P. Johnson, Jr., P. Wu, J. Li, M. Coons, C. Tam, B. Brillantes, H. Sampang, K. Mortara, K. K. Bowman, K. R. Clark, A. Estevez, Z. Xie, H. Verschoof, M. Grimwood, C. Dehnhardt, J. C. Andrez, T. Focken, D. P. Sutherlin, B. S. Safina, M. A. Starovasnik, D. F. Ortwine, Y. Franke, C. J. Cohen, D. H. Hackos, C. M. Koth and J. Payandeh (2015). “Structural basis of Nav1.7 inhibition by an isoform-selective small-molecule antagonist.” Science 350(6267): aac5464.

An, W. F., M. R. Bowlby, M. Betty, J. Cao, H. P. Ling, G. Mendoza, J. W. Hinson, K. I. Mattsson, B. W. Strassle, J. S. Trimmer and K. J. Rhodes (2000). “Modulation of A-type potassium channels by a family of calcium sensors.” Nature 403(6769): 553–556.

Antonucci, D. E., S. T. Lim, S. Vassanelli and J. S. Trimmer (2001). “Dynamic localization and clustering of dendritic Kv2.1 voltage-dependent potassium channels in developing hippocampal neurons.” Neuroscience 108(1): 69–81.

Armstrong, C. M. and F. Bezanilla (1973). “Currents related to movement of the gating particles of the sodium channels.” Nature 242(5398): 459–461.

Benndorf, K., R. Koopmann, C. Lorra and O. Pongs (1994). “Gating and conductance properties of a human delayed rectifier K+ channel expressed in frog oocytes.” J Physiol 477(Pt 1): 1–14.

Bezanilla, F. (2008). “How membrane proteins sense voltage.” Nat Rev Mol Cell Biol 9(4): 323–332.

Bezanilla, F. (2018). “Gating currents.” J Gen Physiol 150(7): 911–932.

Bishop, H. I., D. Guan, E. Bocksteins, L. K. Parajuli, K. D. Murray, M. M. Cobb, H. Misonou, K. Zito, R. C. Foehring and J. S. Trimmer (2015). “Distinct Cell- and Layer-Specific Expression Patterns and Independent Regulation of Kv2 Channel Subtypes in Cortical Pyramidal Neurons.” J Neurosci 35(44): 14922–14942.

Bocksteins, E. (2016). “Kv5, Kv6, Kv8, and Kv9 subunits: No simple silent bystanders.” J Gen Physiol 147(2): 105–125.

Bolte, S. and F. P. Cordelieres (2006). “A guided tour into subcellular colocalization analysis in light microscopy.” J Microsc 224(Pt 3): 213–232.

Catterall, W. A., S. Cestele, V. Yarov-Yarovoy, F. H. Yu, K. Konoki and T. Scheuer (2007). “Voltage-gated ion channels and gating modifier toxins.” Toxicon 49(2): 124–141.

Cerda, O. and J. S. Trimmer (2011). “Activity-dependent phosphorylation of neuronal Kv2.1 potassium channels by CDK5.” J Biol Chem 286(33): 28738–28748.

Dai, X. Q., J. Kolic, P. Marchi, S. Sipione and P. E. Macdonald (2009). “SUMOylation regulates Kv2.1 and modulates pancreatic beta-cell excitability.” J Cell Sci 122(Pt 6): 775–779.

Dai, X. Q., J. E. Manning Fox, D. Chikvashvili, M. Casimir, G. Plummer, C. Hajmrle, A. F. Spigelman, T. Kin, D. Singer-Lahat, Y. Kang, A. M. Shapiro, H. Y. Gaisano, I. Lotan and P. E. Macdonald (2012). “The voltage-dependent potassium channel subunit Kv2.1 regulates insulin secretion from rodent and human islets independently of its electrical function.” Diabetologia 55(6): 1709–1720.

Deutsch, E., A. V. Weigel, E. J. Akin, P. Fox, G. Hansen, C. J. Haberkorn, R. Loftus, D. Krapf and M. M. Tamkun (2012). “Kv2.1 cell surface clusters are insertion platforms for ion channel delivery to the plasma membrane.” Mol Biol Cell 23(15): 2917–2929.

Dockendorff, C., D. M. Gandhi, I. H. Kimball, K. S. Eum, R. Rusinova, H. I. Ingolfsson, R. Kapoor, T. Peyear, M. W. Dodge, S. F. Martin, R. W. Aldrich, O. S. Andersen and J. T. Sack (2018). “Synthetic Analogues of the Snail Toxin 6-Bromo-2-mercaptotryptamine Dimer (BrMT) Reveal That Lipid Bilayer Perturbation Does Not Underlie Its Modulation of Voltage-Gated Potassium Channels.” Biochemistry 57(18): 2733–2743.

Du, J., J. H. Tao-Cheng, P. Zerfas and C. J. McBain (1998). “The K+ channel, Kv2.1, is apposed to astrocytic processes and is associated with inhibitory postsynaptic membranes in hippocampal and cortical principal neurons and inhibitory interneurons.” Neuroscience 84(1): 37–48.

Dunn, K. W., M. M. Kamocka and J. H. McDonald (2011). “A practical guide to evaluating colocalization in biological microscopy.” American Journal of Physiology-Cell Physiology 300(4): C723–C742.

Eichel, K. and M. von Zastrow (2018). “Subcellular Organization of GPCR Signaling.” Trends Pharmacol Sci 39(2): 200–208.

Feinshreiber, L., D. Singer-Lahat, R. Friedrich, U. Matti, A. Sheinin, O. Yizhar, R. Nachman, D. Chikvashvili, J. Rettig, U. Ashery and I. Lotan (2010). “Non-conducting function of the Kv2.1 channel enables it to recruit vesicles for release in neuroendocrine and nerve cells.” J Cell Sci 123(Pt 11): 1940–1947.

Fletcher-Taylor, S., P. Thapa, R. J. Sepela, R. Kaakati, V. Yarov-Yarovoy, J. T. Sack and B. E. Cohen (2020). “Distinguishing Potassium Channel Resting State Conformations in Live Cells with Environment-Sensitive Fluorescence.” ACS Chem Neurosci.

Fox, P. D., C. J. Haberkorn, E. J. Akin, P. J. Seel, D. Krapf and M. M. Tamkun (2015). “Induction of stable ER-plasma-membrane junctions by Kv2.1 potassium channels.” J Cell Sci 128(11): 2096–2105.

Fox, P. D., C. J. Haberkorn, A. V. Weigel, J. L. Higgins, E. J. Akin, M. J. Kennedy, D. Krapf and M. M. Tamkun (2013). “Plasma membrane domains enriched in cortical endoplasmic reticulum function as membrane protein trafficking hubs.” Mol Biol Cell 24(17): 2703–2713.

Fox, P. D., R. J. Loftus and M. M. Tamkun (2013). “Regulation of Kv2.1 K(+) conductance by cell surface channel density.” J Neurosci 33(3): 1259–1270.

Frech, G. C., A. M. VanDongen, G. Schuster, A. M. Brown and R. H. Joho (1989). “A novel potassium channel with delayed rectifier properties isolated from rat brain by expression cloning.” Nature 340(6235): 642–645.

Gamper, N., J. D. Stockand and M. S. Shapiro (2005). “The use of Chinese hamster ovary (CHO) cells in the study of ion channels.” J Pharmacol Toxicol Methods 51(3): 177–185.

Gayet-Primo, J., D. B. Yaeger, R. A. Khanjian and T. Puthussery (2018). “Heteromeric KV2/KV8.2 Channels Mediate Delayed Rectifier Potassium Currents in Primate Photoreceptors.” J Neurosci 38(14): 3414–3427.

Gordon, E., T. K. Roepke and G. W. Abbott (2006). “Endogenous KCNE subunits govern Kv2.1 K+ channel activation kinetics in Xenopus oocyte studies.” Biophys J 90(4): 1223–1231.

Gupta, K., M. Zamanian, C. Bae, M. Milescu, D. Krepkiy, D. C. Tilley, J. T. Sack, V. Yarov-Yarovoy, J. I. Kim and K. J. Swartz (2015). “Tarantula toxins use common surfaces for interacting with Kv and ASIC ion channels.” Elife 4: e06774.

Herrington, J., Y. P. Zhou, R. M. Bugianesi, P. M. Dulski, Y. Feng, V. A. Warren, M. M. Smith, M. G. Kohler, V. M. Garsky, M. Sanchez, M. Wagner, K. Raphaelli, P. Banerjee, C. Ahaghotu, D. Wunderler, B. T. Priest, J. T. Mehl, M. L. Garcia, O. B. McManus, G. J. Kaczorowski and R. S. Slaughter (2006). “Blockers of the delayed-rectifier potassium current in pancreatic beta-cells enhance glucose-dependent insulin secretion.” Diabetes 55(4): 1034–1042.

Irannejad, R., J. C. Tomshine, J. R. Tomshine, M. Chevalier, J. P. Mahoney, J. Steyaert, S. G. Rasmussen, R. K. Sunahara, H. El-Samad, B. Huang and M. von Zastrow (2013). “Conformational biosensors reveal GPCR signalling from endosomes.” Nature 495(7442): 534–538.

Islas, L. D. and F. J. Sigworth (1999). “Voltage sensitivity and gating charge in Shaker and Shab family potassium channels.” J Gen Physiol 114(5): 723–742.

Jara-Oseguera, A., I. G. Ishida, G. E. Rangel-Yescas, N. Espinosa-Jalapa, J. A. Perez-Guzman, D. Elias-Vinas, R. Le Lagadec, T. Rosenbaum and L. D. Islas (2011). “Uncoupling charge movement from channel opening in voltage-gated potassium channels by ruthenium complexes.” J Biol Chem 286(18): 16414–16425.

Johnson, B., A. N. Leek, L. Sole, E. E. Maverick, T. P. Levine and M. M. Tamkun (2018). “Kv2 potassium channels form endoplasmic reticulum/plasma membrane junctions via interaction with VAPA and VAPB.” Proc Natl Acad Sci U S A 115(31): E7331–e7340.

Kaczmarek, L. K. (2006). “Non-conducting functions of voltage-gated ion channels.” Nat Rev Neurosci 7(10): 761–771.

Kang, S. K., C. G. Vanoye, S. N. Misra, D. M. Echevarria, J. D. Calhoun, J. B. O’Connor, K. L. Fabre, D. McKnight, L. Demmer, P. Goldenberg, L. E. Grote, I. Thiffault, C. Saunders, K. A. Strauss, A. Torkamani, J. van der Smagt, K. van Gassen, R. P. Carson, J. Diaz, E. Leon, J. E. Jacher, M. C. Hannibal, J. Litwin, N. R. Friedman, A. Schreiber, B. Lynch, A. Poduri, E. D. Marsh, E. M. Goldberg, J. J. Millichap, A. L. George Jr and J. A. Kearney (2019). “Spectrum of KV2.1 Dysfunction in KCNB1-Associated Neurodevelopmental Disorders.” Annals of Neurology 86(6): 899–912.

Kihira, Y., T. O. Hermanstyne and H. Misonou (2010). “Formation of heteromeric Kv2 channels in mammalian brain neurons.” J Biol Chem 285(20): 15048–15055.

Kimm, T., Z. M. Khaliq and B. P. Bean (2015). “Differential Regulation of Action Potential Shape and Burst-Frequency Firing by BK and Kv2 Channels in Substantia Nigra Dopaminergic Neurons.” J Neurosci 35(50): 16404–16417.

Kirmiz, M., T. E. Gillies, E. J. Dickson and J. S. Trimmer (2019). “Neuronal ER-plasma membrane junctions organized by Kv2-VAP pairing recruit Nir proteins and affect phosphoinositide homeostasis.” J Biol Chem 294(47): 17735–17757.

Kirmiz, M., S. Palacio, P. Thapa, A. N. King, J. T. Sack and J. S. Trimmer (2018). “Remodeling neuronal ER-PM junctions is a conserved nonconducting function of Kv2 plasma membrane ion channels.” Mol Biol Cell 29(20): 2410–2432.

Kirmiz, M., N. C. Vierra, S. Palacio and J. S. Trimmer (2018). “Identification of VAPA and VAPB as Kv2 Channel-Interacting Proteins Defining Endoplasmic Reticulum-Plasma Membrane Junctions in Mammalian Brain Neurons.” J Neurosci 38(35): 7562–7584.

Lee, H. C., J. M. Wang and K. J. Swartz (2003). “Interaction between extracellular Hanatoxin and the resting conformation of the voltage-sensor paddle in Kv channels.” Neuron 40(3): 527–536.

Lewis, G. N. (1925). “A New Principle of Equilibrium.” Proc Natl Acad Sci U S A 11(3): 179–183.

Li, H., W. Guo, H. Xu, R. Hood, A. T. Benedict and J. M. Nerbonne (2001). “Functional expression of a GFP-tagged Kv1.5 alpha-subunit in mouse ventricle.” Am J Physiol Heart Circ Physiol 281(5): H1955–1967.

Lin, M. Z. and M. J. Schnitzer (2016). “Genetically encoded indicators of neuronal activity.” Nat Neurosci 19(9): 1142–1153.

Liu, P. W. and B. P. Bean (2014). “Kv2 channel regulation of action potential repolarization and firing patterns in superior cervical ganglion neurons and hippocampal CA1 pyramidal neurons.” J Neurosci 34(14): 4991–5002.

Long, S. B., E. B. Campbell and R. Mackinnon (2005). “Voltage sensor of Kv1.2: structural basis of electromechanical coupling.” Science 309(5736): 903–908.

Long, S. B., X. Tao, E. B. Campbell and R. MacKinnon (2007). “Atomic structure of a voltage-dependent K+ channel in a lipid membrane-like environment.” Nature 450(7168): 376–382.

MacDonald, P. E., A. M. Salapatek and M. B. Wheeler (2003). “Temperature and redox state dependence of native Kv2.1 currents in rat pancreatic beta-cells.” J Physiol 546(Pt 3): 647–653.

Maletic-Savatic, M., N. J. Lenn and J. S. Trimmer (1995). “Differential spatiotemporal expression of K+ channel polypeptides in rat hippocampal neurons developing in situ and in vitro.” J Neurosci 15(5 Pt 2): 3840–3851.

McCord, M. C. and E. Aizenman (2013). “Convergent Ca2+ and Zn2+ signaling regulates apoptotic Kv2.1 K+ currents.” Proc Natl Acad Sci U S A 110(34): 13988–13993.

McCormack, K., S. Santos, M. L. Chapman, D. S. Krafte, B. E. Marron, C. W. West, M. J. Krambis, B. M. Antonio, S. G. Zellmer, D. Printzenhoff, K. M. Padilla, Z. Lin, P. K. Wagoner, N. A. Swain, P. A. Stupple, M. de Groot, R. P. Butt and N. A. Castle (2013). “Voltage sensor interaction site for selective small molecule inhibitors of voltage-gated sodium channels.” Proc Natl Acad Sci U S A 110(29): E2724–2732.

McCrossan, Z. A., T. K. Roepke, A. Lewis, G. Panaghie and G. W. Abbott (2009). “Regulation of the Kv2.1 Potassium Channel by MinK and MiRP1.” Journal of Membrane Biology 228(1): 1–14.

McDonough, S. I., I. M. Mintz and B. P. Bean (1997). “Alteration of P-type calcium channel gating by the spider toxin omega-Aga-IVA.” Biophys J 72(5): 2117–2128.

Milescu, M., F. Bosmans, S. Lee, A. A. Alabi, J. I. Kim and K. J. Swartz (2009). “Interactions between lipids and voltage sensor paddles detected with tarantula toxins.” Nat Struct Mol Biol 16(10): 1080–1085.

Misonou, H., D. P. Mohapatra, M. Menegola and J. S. Trimmer (2005). “Calcium- and metabolic state-dependent modulation of the voltage-dependent Kv2.1 channel regulates neuronal excitability in response to ischemia.” J Neurosci 25(48): 11184–11193.

Misonou, H., D. P. Mohapatra, E. W. Park, V. Leung, D. Zhen, K. Misonou, A. E. Anderson and J. S. Trimmer (2004). “Regulation of ion channel localization and phosphorylation by neuronal activity.” Nat Neurosci 7(7): 711–718.

Murakoshi, H., G. Shi, R. H. Scannevin and J. S. Trimmer (1997). “Phosphorylation of the Kv2.1 K^+^ Channel Alters Voltage-Dependent Activation.” Molecular Pharmacology 52(5): 821–828.

O’Connell, K. M., R. Loftus and M. M. Tamkun (2010). “Localization-dependent activity of the Kv2.1 delayed-rectifier K+ channel.” Proc Natl Acad Sci U S A 107(27): 12351–12356.

Opitz-Araya, X. and A. Barria (2011). “Organotypic hippocampal slice cultures.” J Vis Exp(48).

Park, K. S., D. P. Mohapatra, H. Misonou and J. S. Trimmer (2006). “Graded regulation of the Kv2.1 potassium channel by variable phosphorylation.” Science 313(5789): 976–979.

Peltola, M. A., J. Kuja-Panula, S. E. Lauri, T. Taira and H. Rauvala (2011). “AMIGO is an auxiliary subunit of the Kv2.1 potassium channel.” EMBO Rep 12(12): 1293–1299.

Peretz, A., L. Pell, Y. Gofman, Y. Haitin, L. Shamgar, E. Patrich, P. Kornilov, O. Gourgy-Hacohen, N. Ben-Tal and B. Attali (2010). “Targeting the voltage sensor of Kv7.2 voltage-gated K+ channels with a new gating-modifier.” Proc Natl Acad Sci U S A 107(35): 15637–15642.

Plant, L. D., E. J. Dowdell, I. S. Dementieva, J. D. Marks and S. A. Goldstein (2011). “SUMO modification of cell surface Kv2.1 potassium channels regulates the activity of rat hippocampal neurons.” J Gen Physiol 137(5): 441–454.

Pologruto, T. A., B. L. Sabatini and K. Svoboda (2003). “ScanImage: flexible software for operating laser scanning microscopes.” Biomed Eng Online 2: 13.

Ramu, Y., Y. Xu and Z. Lu (2006). “Enzymatic activation of voltage-gated potassium channels.” Nature 442(7103): 696–699.

Sack, J. T., R. W. Aldrich and W. F. Gilly (2004). “A gastropod toxin selectively slows early transitions in the Shaker K channel’s activation pathway.” J Gen Physiol 123(6): 685–696.

Sack, J. T., N. Stephanopoulos, D. C. Austin, M. B. Francis and J. S. Trimmer (2013). “Antibody-guided photoablation of voltage-gated potassium currents.” J Gen Physiol 142(3): 315–324.

Schmalhofer, W. A., J. Calhoun, R. Burrows, T. Bailey, M. G. Kohler, A. B. Weinglass, G. J. Kaczorowski, M. L. Garcia, M. Koltzenburg and B. T. Priest (2008). “ProTx-II, a selective inhibitor of NaV1.7 sodium channels, blocks action potential propagation in nociceptors.” Mol Pharmacol 74(5): 1476–1484.

Schneider, C. A., W. S. Rasband and K. W. Eliceiri (2012). “NIH Image to ImageJ: 25 years of image analysis.” Nat Methods 9(7): 671–675.

Schneider, M. F. and W. K. Chandler (1973). “Voltage dependent charge movement of skeletal muscle: a possible step in excitation-contraction coupling.” Nature 242(5395): 244–246.

Scholle, A., S. Dugarmaa, T. Zimmer, M. Leonhardt, R. Koopmann, B. Engeland, O. Pongs and J. Benndorf (2004). “Rate-limiting reactions determining different activation kinetics of Kv1.2 and Kv2.1 channels.” J Membr Biol 198(2): 103–112.

Seoh, S. A., D. Sigg, D. M. Papazian and F. Bezanilla (1996). “Voltage-sensing residues in the S2 and S4 segments of the Shaker K+ channel.” Neuron 16(6): 1159–1167.

Shi, G., K. Nakahira, S. Hammond, K. J. Rhodes, L. E. Schechter and J. S. Trimmer (1996). “Beta subunits promote K+ channel surface expression through effects early in biosynthesis.” Neuron 16(4): 843–852.

Shibata, R., H. Misonou, C. R. Campomanes, A. E. Anderson, L. A. Schrader, L. C. Doliveira, K. I. Carroll, J. D. Sweatt, K. J. Rhodes and J. S. Trimmer (2003). “A fundamental role for KChIPs in determining the molecular properties and trafficking of Kv4.2 potassium channels.” J Biol Chem 278(38): 36445–36454.

Singer-Lahat, D., A. Sheinin, D. Chikvashvili, S. Tsuk, D. Greitzer, R. Friedrich, L. Feinshreiber, U. Ashery, M. Benveniste, E. S. Levitan and I. Lotan (2007). “K+ channel facilitation of exocytosis by dynamic interaction with syntaxin.” J Neurosci 27(7): 1651–1658.

Speca, D. J., G. Ogata, D. Mandikian, H. I. Bishop, S. W. Wiler, K. Eum, H. J. Wenzel, E. T. Doisy, L. Matt, K. L. Campi, M. S. Golub, J. M. Nerbonne, J. W. Hell, B. C. Trainor, J. T. Sack, P. A. Schwartzkroin and J. S. Trimmer (2014). “Deletion of the Kv2.1 delayed rectifier potassium channel leads to neuronal and behavioral hyperexcitability.” Genes Brain Behav 13(4): 394–408.

Stoppini, L., P. A. Buchs and D. Muller (1991). “A simple method for organotypic cultures of nervous tissue.” J Neurosci Methods 37(2): 173–182.

Swartz, K. J. (2007). “Tarantula toxins interacting with voltage sensors in potassium channels.” Toxicon 49(2): 213–230.

Tanabe, T., K. G. Beam, J. A. Powell and S. Numa (1988). “Restoration of excitation-contraction coupling and slow calcium current in dysgenic muscle by dihydropyridine receptor complementary DNA.” Nature 336(6195): 134–139.

Tao, X., A. Lee, W. Limapichat, D. A. Dougherty and R. MacKinnon (2010). “A gating charge transfer center in voltage sensors.” Science 328(5974): 67–73.

Tilley, D. C., J. M. Angueyra, K. S. Eum, H. Kim, L. H. Chao, A. W. Peng and J. T. Sack (2019). “The tarantula toxin GxTx detains K(+) channel gating charges in their resting conformation.” J Gen Physiol 151(3): 292–315.

Tilley, D. C., K. S. Eum, S. Fletcher-Taylor, D. C. Austin, C. Dupre, L. A. Patron, R. L. Garcia, K. Lam, V. Yarov-Yarovoy, B. E. Cohen and J. T. Sack (2014). “Chemoselective tarantula toxins report voltage activation of wild-type ion channels in live cells.” Proc Natl Acad Sci U S A 111(44): E4789–4796.

Trapani, J. G. and S. J. Korn (2003). “Control of ion channel expression for patch clamp recordings using an inducible expression system in mammalian cell lines.” BMC Neurosci 4: 15.

Trimmer, J. S. (1991). “Immunological identification and characterization of a delayed rectifier K+ channel polypeptide in rat brain.” Proc Natl Acad Sci U S A 88(23): 10764–10768.

Tsvetanova, N. G., R. Irannejad and M. von Zastrow (2015). “G protein-coupled receptor (GPCR) signaling via heterotrimeric G proteins from endosomes.” J Biol Chem 290(11): 6689–6696.

Turner, M., D. E. Anderson, P. Bartels, M. Nieves-Cintron, A. M. Coleman, P. B. Henderson, K. N. M. Man, P. Y. Tseng, V. Yarov-Yarovoy, D. M. Bers, M. F. Navedo, M. C. Horne, J. B. Ames and J. W. Hell (2020). “α-Actinin-1 promotes activity of the L-type Ca(2+) channel Ca(v) 1.2.” Embo j 39(5): e102622.

Vacher, H., D. P. Mohapatra and J. S. Trimmer (2008). “Localization and targeting of voltage-dependent ion channels in mammalian central neurons.” Physiol Rev 88(4): 1407–1447.

Vierra, N. C., M. Kirmiz, D. van der List, L. F. Santana and J. S. Trimmer (2019). “Kv2.1 mediates spatial and functional coupling of L-type calcium channels and ryanodine receptors in mammalian neurons.” Elife 8.

Weigel, A. V., P. D. Fox, E. J. Akin, K. H. Ecklund, M. M. Tamkun and D. Krapf (2012). “Size of cell-surface Kv2.1 domains is governed by growth fluctuations.” Biophys J 103(8): 1727–1734.

Weigel, A. V., M. M. Tamkun and D. Krapf (2013). “Quantifying the dynamic interactions between a clathrin-coated pit and cargo molecules.” Proceedings of the National Academy of Sciences 110(48): E4591–E4600.

Woods, G. and K. Zito (2008). “Preparation of gene gun bullets and biolistic transfection of neurons in slice culture.” J Vis Exp(12).

Xu, H., T. Li, A. Rohou, C. P. Arthur, F. Tzakoniati, E. Wong, A. Estevez, C. Kugel, Y. Franke, J. Chen, C. Ciferri, D. H. Hackos, C. M. Koth and J. Payandeh (2019). “Structural Basis of Nav1.7 Inhibition by a Gating-Modifier Spider Toxin.” Cell 176(5): 1238–1239.

Yang, F. and J. Zheng (2014). “High temperature sensitivity is intrinsic to voltage-gated potassium channels.” Elife 3: e03255.

Yang, W. and R. Yuste (2017). “In vivo imaging of neural activity.” Nat Methods 14(4): 349–359.

Zagotta, W. N., T. Hoshi, J. Dittman and R. W. Aldrich (1994). “Shaker potassium channel gating. II: Transitions in the activation pathway.” J Gen Physiol 103(2): 279–319.

Zhang, A. H., G. Sharma, E. A. B. Undheim, X. Jia and M. Mobli (2018). “A complicated complex: Ion channels, voltage sensing, cell membranes and peptide inhibitors.” Neurosci Lett 679: 35–47.

Zhang, G., S. Zheng, H. Liu and P. R. Chen (2015). “Illuminating biological processes through site-specific protein labeling.” Chem Soc Rev 44(11): 3405–3417.

Zito, K., G. Knott, G. M. Shepherd, S. Shenolikar and K. Svoboda (2004). “Induction of spine growth and synapse formation by regulation of the spine actin cytoskeleton.” Neuron 44(2): 321–334.

